# A fluorescence-based genetic screen reveals diverse mechanisms silencing small RNA signaling in *E. coli*

**DOI:** 10.1101/2021.05.11.443692

**Authors:** Jiandong Chen, Leann To, Francois de Mets, Xing Luo, Nadim Majdalani, Chin-Hsien Tai, Susan Gottesman

## Abstract

As key players of gene regulation in many bacteria, small regulatory RNAs (sRNAs) associated with the RNA chaperone Hfq shape numerous phenotypic traits, including metabolism, stress response and adaptation as well as virulence. sRNAs can alter target mRNA translation and stability via base pairing. sRNA synthesis is generally under tight transcriptional regulation, but other levels of regulation of sRNA signaling are less well understood. Here we used a fluorescence-based functional screen to identify new regulators that can quench sRNA signaling of the iron-responsive sRNA RyhB in *E. coli*. The identified regulators fell into two classes, general regulators (affecting signaling by many sRNAs) and RyhB-specific regulators; we focused on the specific ones here. General regulators include three Hfq-interacting sRNAs, CyaR, ChiX and McaS, previously found to act through Hfq competition, RNase T, a 3′-5′ exonuclease not previously implicated in sRNA degradation, and YhbS, a putative GCN5-related N-acetyltransferase (GNAT). Two new specific regulators were identified. AspX, a novel 3′end-derived small RNA, specifically represses RyhB signaling via an RNA sponging mechanism. YicC, a previously uncharacterized but widely conserved protein, triggers rapid RyhB degradation via collaboration with the exoribonuclease PNPase. These findings greatly expand our knowledge of regulation of bacterial sRNA signaling and suggest complex regulatory networks for controlling iron homeostasis in bacteria. The fluorescence-based genetic screen system described here is a powerful tool expected to accelerate the discovery of novel regulators of sRNA signaling in many bacteria.

**Significance:** The ability to promptly switch genes on and off allows bacteria to adapt rapidly to changing environments for better survival. Many sRNAs, including RyhB, a sRNA made in response to iron starvation, are important switches in bacteria. We discovered new factors that can keep the sRNA switch off by using a facile genetic screen platform. These new factors include a new RNA sponge and an adaptor protein for a ribonuclease, providing new perspectives on controlling sRNA signaling in bacteria.

## Introduction

Bacterial small regulatory RNAs (sRNAs) are important posttranscriptional regulators of gene expression. Under stress conditions, sRNAs can be rapidly generated to reprogram gene expression, allowing bacteria fast adaptation to environmental or intracellular perturbations [reviewed in (1, 2)]. sRNAs can affect translation or stability of mRNA targets through base-pairing mechanisms, and in many bacteria, the RNA chaperone Hfq is required for this process. Hfq forms a ring-shaped homohexamer and preferentially binds the polyU tail of sRNAs and AAN motifs of mRNAs, using its proximal and distal surface respectively (3). Hfq binding usually protects sRNAs from ribonuclease degradation, and also promotes formation of an RNA duplex between sRNA and cognate mRNAs. Research, mostly from studies in *E. coli* and *Salmonella*, has established a general working model where newly transcribed sRNAs are bound by Hfq pair with mRNAs. The base-pairing is frequently adjacent to the translation initiation region, thus altering ribosome occupancy and translation, and/or recruiting the RNA degradosome to degrade mRNAs [reviewed in (4, 5)]. A number of deviations from this canonical model have been reported as well (4). Although the core components (*e*.*g*. Hfq and sRNAs) and their molecular mechanisms of action have been extensively studied, auxiliary factors that modulate robustness or specificity of sRNAs are less known.

Regulation of sRNAs can occur at multiple levels, including sRNA transcription, RNA stability and effects on the ability of the sRNA to carry out regulation. Although sRNAs are known to be primarily switched on/off through transcriptional control in response to environmental cues, emerging examples suggest that they can be regulated posttranscriptionally as well [reviewed in (6, 7)]. Recent RNA-seq-based approaches (8-11) have identified new RNA transcripts that can titrate sRNAs from their mRNA targets, thus acting as RNA sponges for specific sRNAs. Such sponge RNAs are likely widely present, but only a few have been discovered and studied in detail. Interestingly, in the parallel metazoan microRNA system, posttranscriptional regulation of microRNA pathways is pervasive and contributes significantly to microRNA functions (12, 13). Whether parallel regulatory mechanisms are present in bacterial sRNA pathways remains unknown.

Functional genetic screens provide a powerful discovery tool that allows identification of regulators of sRNA signaling beyond RNA sponges. In this study, we aimed to identify genetic factors that can negatively regulate sRNA signaling in *E. coli*. By combining our sensitive fluorescence-based reporter system that can monitor endogenous sRNA activities in live *E*.*coli* cells with a genomic library screen approach, we identified known and new quenchers of sRNA signaling, including Hfq-dependent sRNAs, a new RNA sponge, a ribonuclease and two conserved proteins of unknown function. Study of the molecular action of the individual regulators revealed that diverse posttranscriptional regulatory mechanisms can silence sRNA signaling in *E. coli*.

## Results and Discussion

### A chromosomal tandem fluorescence reporter system allows monitoring sRNA activities in live *E. coli* cells

Gene reporters are invaluable tools for facile measurement of gene expression. As key posttranscriptional modulators of bacterial gene expression, sRNA reporters, including plasmid-based fluorescent reporters, have been successfully developed to monitor sRNA activities in gram-negative or gram-positive bacteria (14, 15). Since most sRNAs affect translation, reporters that are translationally fused to the target gene are commonly used. However, target mRNA stability is also frequently affected, either directly or indirectly, via the effects on translation (16). Here we devised a chromosome-encoded, tandem fluorescence reporter system for monitoring sRNA activities via fluorescence readout of both the translation and stability of mRNA targets. This mini-operon cassette consists of a gene of interest (GOI) that is translationally fused to the gene encoding red fluorescent protein mCherry; the second gene in the operon encodes superfolder green fluorescent protein sfGFP as an indirect readout for mRNA levels. Transcription of the whole cassette is driven by a constitutive synthetic promoter Cp26 (17) and terminated through a ribosomal T1 terminator (**Fig. S1A**). The whole cassette was inserted at the *lacZ* locus of the *E. coli* genome. We first tested the response of this reporter system to two endogenous sRNAs. We generated translational fusions to mCherry of two genes, *chiP* and *sodB*, both of which are well characterized mRNA targets, repressed by ChiX and RyhB sRNA respectively (18, 19) (**Fig. 1, A and D**), and compared fluorescence signals of cells with or without the corresponding sRNA. In *chiP* reporter cells, endogenous ChiX sRNA is constitutively expressed and strongly inhibits *chiP* expression, resulting in nonfluorescent cells. Deletion of *chiX* led to highly fluorescent cells for both mCherry and GFP (**Fig. 1B**), consistent with the repressive roles of ChiX in inhibiting translation and promoting decay of *chiP* mRNA (18). Quantification of fluorescence for cells grown in liquid culture showed a 30-and 12-fold increase of mCherry and GFP signals respectively in *chiX* null cells (**Fig. 1C**). For the *sodB* reporter, WT cells express low levels of RyhB due to transcriptional repression by Fur and thus show high fluorescence on LB agar (19). Deletion of *fur* relieved repression (**Fig. S1B**) and significantly diminished mCherry and GFP fluorescence (**Fig. 1E and F**). This is consistent with previous studies demonstrating that RyhB inhibits translation and promotes decay of the *sodB* mRNA (16). Such changes in fluorescence resulted from RyhB repression of *sodB*, as mutating either *ryhB* or the RNA chaperone gene *hfq* fully restored *sodB* reporter fluorescence, for both mCherry and GFP. These results demonstrated that our tandem fluorescence reporter system can faithfully monitor endogenous sRNA activities in live *E*.*coli*, via fluorescence signals, both qualitatively and quantitatively.

**Figure 1.**
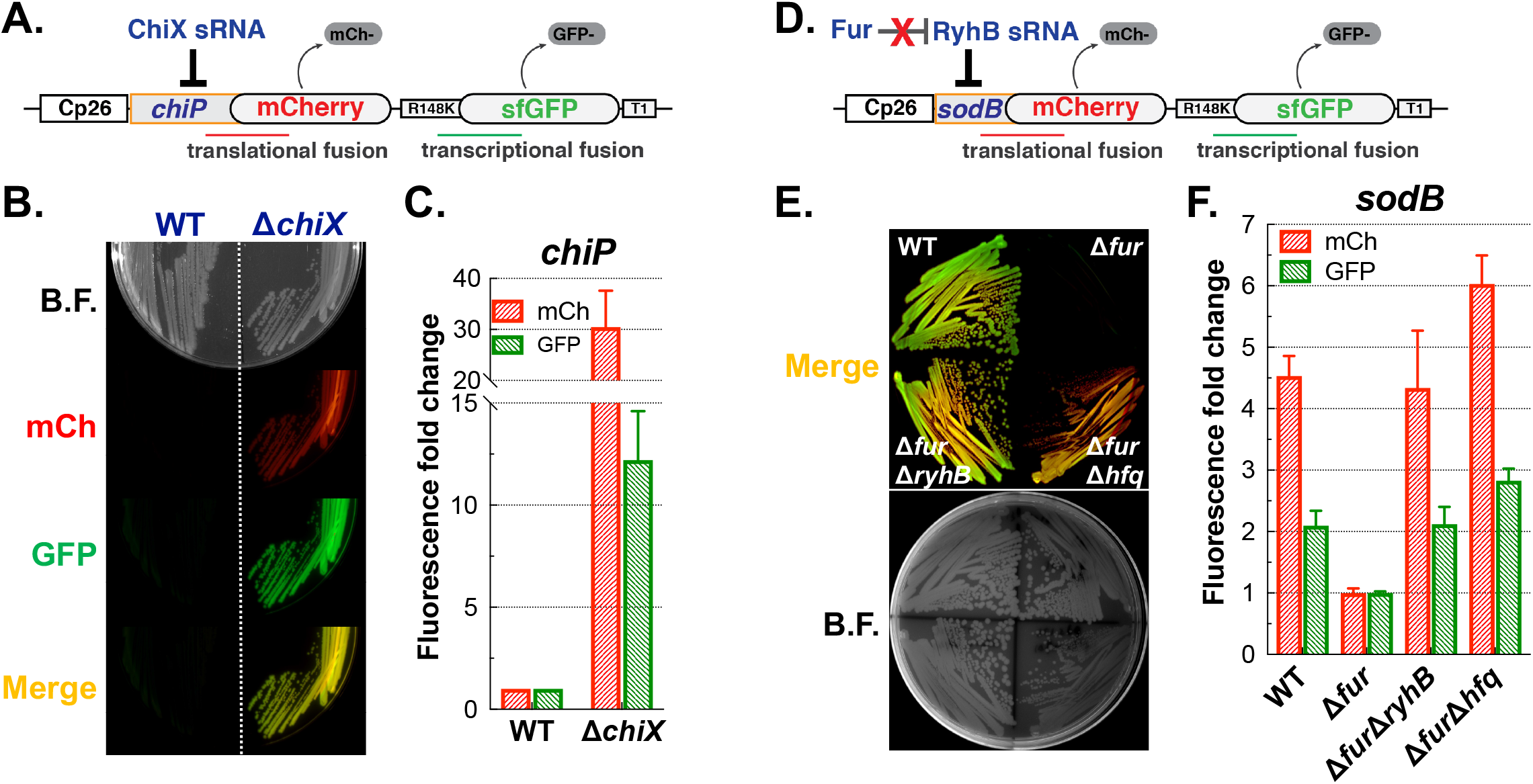
A chromosomal tandem fluorescence reporter system allows facile monitoring of endogenous sRNA activity in *E. coli*. Schematic of the *chiP* (**A**) or *sodB* (**D**) dual-fluorescence reporter. Colony fluorescence on LB agar for reporter strains carrying translational fusions to *chiP* (**B**, WT (JC1200); Δ*chiX* (JC1244)) or *sodB* (**E**, WT (JC1246); Δ*fur* (JC1248); Δ*fur*Δ*ryhB* (JC1252); Δ*fur*Δ*hfq* (JC1269)) was imaged using the Bio-Rad ChemiDoc MP Imaging System. Bright field imaging (B.F.) was used to visualize all colonies on the plate. *chiP* (**C**) or *sodB* (**F**) reporter fluorescence from cultures grown in LB for 6 h was measured using the TECAN Spark 10M microplate reader. Representative images for colony fluorescence are shown. For fluorescence quantification, 3-6 individual clones of each strain were grown and measured in 96-well microplates; data are plotted as mean and standard deviation. A detailed description of fluorescence measurement and quantification is included in Materials and Methods.

Besides the two mRNAs described above, we also measured both positive and negative sRNA/Hfq-mediated gene regulation of other mRNA targets, including *rpoS, mutS, ompF* and *ompX*, by comparing expression of appropriate fusions in WT and Δ*hfq* cells (**Fig. S1C**). Hfq positively regulates *rpoS* expression via three sRNAs (20), and deletion of *hfq* indeed decreased RpoS-mCherry fluorescence by 57%. *mutS* is repressed by Hfq, both with and without sRNAs (21), and MutS-mCherry fluorescence increased 4-fold in the *hfq* mutant (**Fig. S1C**). *ompF* and *ompX* are repressed by multiple sRNAs (22-24) and deletion of *hfq* increased mCherry fluorescence by 3-fold and 1.6-fold respectively, consistent with loss of repression from endogenous sRNAs.

The design of our tandem fluorescence reporters made it possible to monitor effects on mRNA abundance in addition to translation. As sRNAs can alter target gene translation and/or stability, our reporter system allows us to independently assess each of these roles for a given sRNA. For most sRNA targets, including *chiP* and *sodB*, sRNA regulation affects both mRNA translation and stability; thus disruption of sRNA function leads to changes of both the translational mCherry fusion and the transcriptional GFP fusion, with greater changes for mCherry (**Figs. 1 and S1C**). For genes primarily regulated at the level of translation by sRNA/Hfq, e.g. *rpoS* and *mutS* (20, 21), deletion of *hfq* significantly changed mCherry fluorescence but had little effect on GFP (**Fig. S1C**). We expect that for mRNAs primarily regulated at the level of stability, mCherry and GFP fluorescence will be similarly affected. Therefore, this reporter can provide rapid insight into the mechanisms of sRNA function in bacteria. As a negative control, we engineered RBS1, a synthetic consensus ribosome-binding sequence, upstream of the mCherry reporter gene. Notably, fluorescence signals were minimally perturbed (1.2-fold increase) by deletion of *hfq* in this control strain, suggesting that Hfq itself does not affect mCherry or GFP translation or maturation.

In addition to monitoring endogenous sRNA/Hfq activities, we also determined the usability of our fluorescent reporter for monitoring sRNA regulation from multicopy plasmids (**Fig. S1D**), a common approach for sRNA research. In agreement with the expected translational activation, overproduction of ArcZ, DsrA or RprA sRNA increased *rpoS*-mCherry expression significantly but had little effect on GFP production. In general, positive regulation may be primarily translational. We have previously found that DsrA overproduction increased translation 120-fold, while mRNA levels increased only 6-fold (25), consistent with this result. For mRNAs regulated for both translation and stability, overproduction of MicF in the *ompF*-mCherry reporter strain or RyhB in the *sodB*-mCherry reporter strain drastically reduced both mCherry and GFP production. As expected, multicopy sRNAs showed much greater effects on reporter fusion expression than endogenous sRNAs. Taken together, these data demonstrate that a fluorescence-based, chromosome-encoded reporter system provides a simple monitor for sRNA-mediated gene regulation in live *E. coli*.

### Fluorescence-based genomic library screen identified genes elevating reporter fusion expression

This facile reporter system allowed us to conduct unbiased genetic screens to identify genetic factors important for sRNA signaling. To identify negative regulators of sRNA signaling, we combined a genomic library approach with the fluorescence-based reporter system (**Fig. 2A**; a detailed screen strategy is described in **Fig. S2**). We transformed an *E. coli* genomic plasmid library consisting of 1.5-5 kb genomic fragments into the *chiP* or *sodB* reporter strain, both of which exhibited low fluorescence (**Fig. 1B: WT and 1E: Δ*fur***), then screened for clones with elevated fluorescence signals on agar plates (500-1000 CFUs per plate). We screened ∼45,000 individual clones for the *chiP* fusion and ∼17,800 for the *sodB* reporter, and identified 25 and 26 clones respectively that had significantly elevated fluorescence signals (**Fig. S3A**). Both mCherry and GFP were affected by the identified regulators, with mCherry showing greater effects; we show only mCherry effects in the rest of this study.

**Figure 2.**
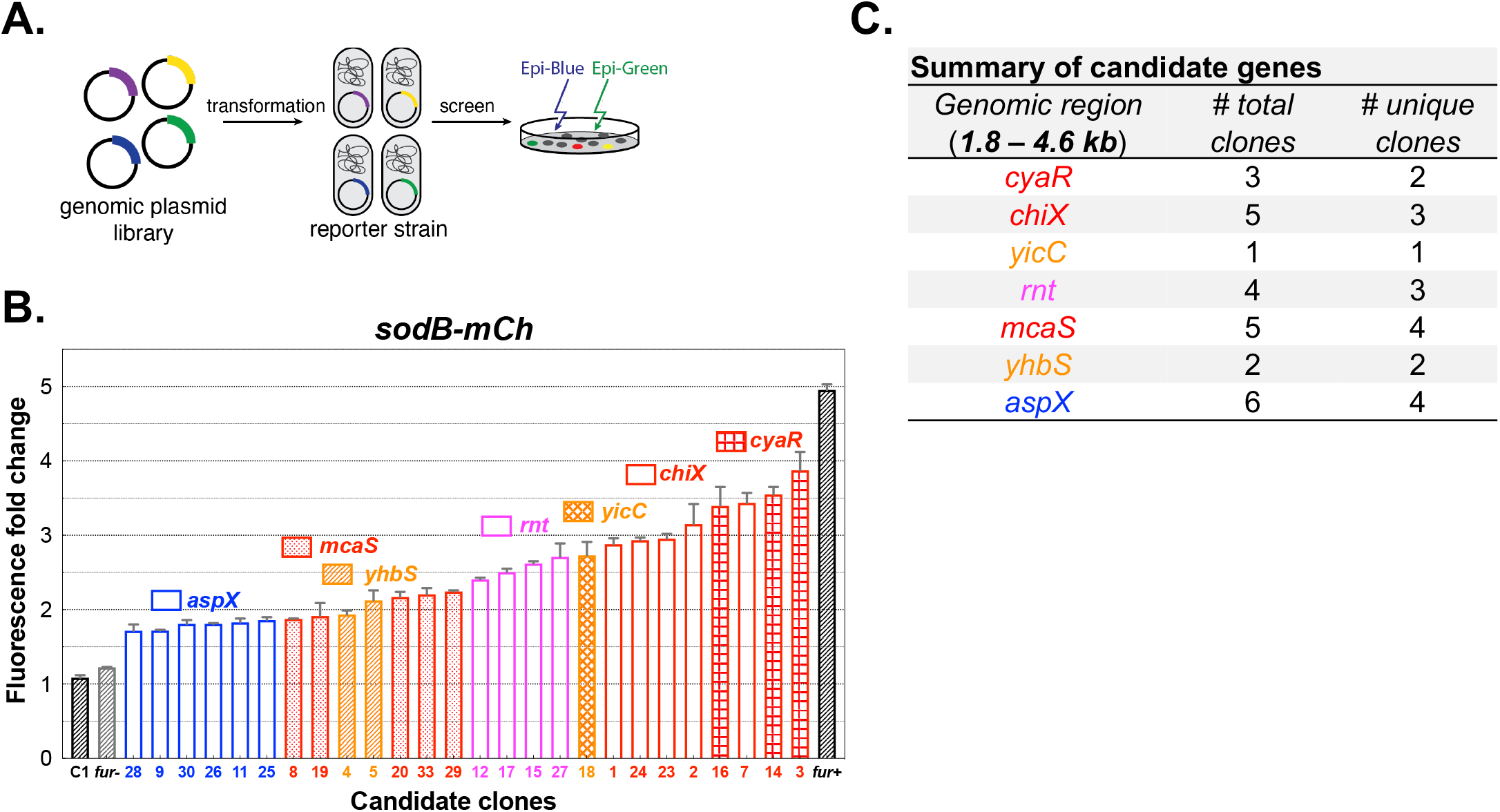
A fluorescence-based genomic library screen identified genomic fragments elevating *sodB* reporter expression. **A**. Schematic of the fluorescence-based genomic library suppressor screen. A plasmid library consisting of 1.5-5 kb *E. coli* genomic fragments was introduced into the *sodB* reporter strain (JC1248). Transformants showing significantly elevated fluorescence were isolated for verification and further analysis. **B**. Relative fluorescence of controls (C1, *fur*- and *fur*+) and candidate clones from the genomic library screen. Cells were grown in 100 μL of LB+Amp in 96-well microplates with shaking (250 rpm) at 37°C for 6 h, and fluorescence signals were measured at the endpoint. Colony fluorescence of the same set of candidate clones are shown in **Fig. S3D**. C1 is a control clone showing basal fluorescence picked from an original screen plate, and *fur*-is the parental reporter strain with the empty vector. Both strains served as basal fluorescence controls. The *fur*+ strain served as a positive control. The original screen identified 33 brighter clones (#1 - #33), and seven were dropped after further examination, due to insignificant increase (<1.5-fold) of fluorescence. **C**. Summary of total and unique clones aligned to unique genomic loci (*cyaR, chiX, yicC, rnt, mcaS, yhbS, aspX*), based on sequencing and mapping of 26 positive clones. Detailed mapping of individual clones to each locus is shown in **Fig. S3E**.

Deleting *chiX* increased ChiP-mCherry fluorescence by 74-fold; the 25 identified positive clones increased *chiP*-mCherry expression by 16-to 95-fold (**Fig. S3B**). Plasmids from eight clones that expressed various levels of mCherry fluorescence were sequenced and all were found to include a genomic region of the *chbBCARFG* operon (**Fig. S3C**). The intergenic region between *chbB* and *chbC* (*chbBC* IGR) encodes an RNA decoy which can base pair with and destabilize ChiX sRNA (18). PCR examination using *chbBC* IGR specific primers for the remaining 17 candidate plasmids found all had the *chbBC* IGR sequence, reinforcing the idea that ChiX sRNA activities are primarily controlled by this RNA decoy under these screening conditions. Our data also supported the feasibility of applying our genetic screen strategy to identify silencers of sRNA signaling. Notably, in this screen only highly fluorescent clones were selected, and thus genetic factors with modest effects would have been missed. As seen below, that includes factors identified with another reporter.

For the *sodB* reporter, our screen identified 26 positive clones, increasing mCherry fluorescence by 1.8-to 3.9-fold (**Figs. 2B and S3D**). Sequencing mapped the inserts to seven different genomic loci (**Figs. 2C and S3E**). In all cases except one (plasmid containing the *yicC* locus), two or more unique clones were identified, and sibling clones were also found in five out of seven groups (**Fig. 2C**). Additionally, clones in the same group largely displayed similar effects on *sodB*-mCherry expression (**Fig. 2B**). Collectively, our genetic screen identified unique genomic fragments that can activate reporter gene expression, presumably via alleviating sRNA-mediated repression. Note that for genes or regions to be identified here, they must be present in the plasmid library (not lethal or next to a gene lethal when overexpressed), and must be expressed at higher levels when present on the plasmid. Therefore, additional regions not identified in this screen may exist. Below we focused on plasmids that abrogated RyhB repression of the *sodB* reporter.

### Individual genetic factors suppress RyhB function

The average size of an insert in our genomic plasmid library was ∼3.0 kb, typically encoding two to four genes. To pinpoint the gene responsible for activating the *sodB* fusion, we aligned all sequences from one genomic area to determine common regions (**Fig. S3E**). We then chose the smallest plasmid from each group, performed on-plasmid deletions of individual candidate genes common to each group and measured SodB-mCherry fluorescence. Deletion of individual candidate genes (*cyaR, chiX, mcaS, rnt, yicC, yhbTS* and *aspX*) from the original plasmids completely abolished activation of *sodB*-mCherry expression (**Fig. 3A**), whereas deletion of other genes, including *rph* or *dinD* on the *yicC* plasmid or *yhbQ* or *yhbP* on the *yhbS* plasmid, had little or no effect (**Fig. S3F**). More importantly, expression of the identified individual factors from an IPTG-inducible heterologous promoter significantly increased *sodB*-mCherry expression, leading to 1.7-to 5.4-fold higher fluorescence signals (**Fig. 3B**), demonstrating that each genetic factor is both necessary (**Fig. 3A**) and sufficient (**Fig. 3B**) to elevate SodB-mCherry production. Finally, in a *ryhB* null strain, overproduction of each factor no longer increased *sodB*-mCherry expression (**Fig. 3C**), demonstrating that the stimulatory effect from each factor is a result of derepression of RyhB-mediated *sodB* silencing, rather than a direct effect on the reporter gene itself.

**Figure 3.**
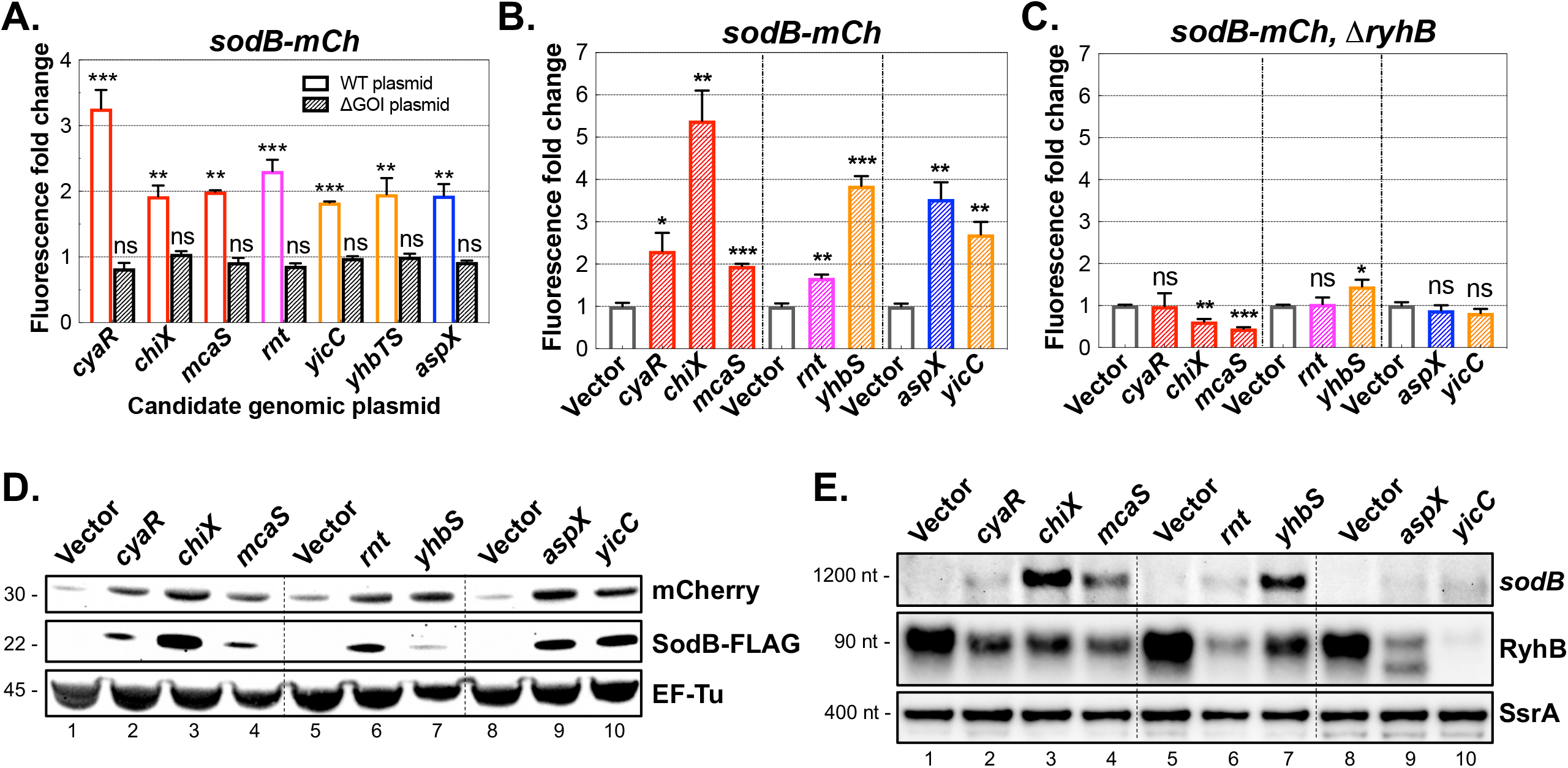
Identification of plasmid-borne regulators necessary and sufficient for suppression of RyhB sRNA function. **A**. mCherry fluorescence of *sodB* reporter strains (JC1248) harboring candidate genomic plasmids (colored bars) or the same plasmids deleted for candidate genes (grey bars). Cells were grown in 96-well microtiter plates with 100 μL of LB+Amp at 37°C for 6 h, and fluorescence was measured at the endpoint, compared to that of the parental reporter strain with the empty vector. Fluorescence of the *sodB* reporter was measured in (**B**) WT (JC1318) or (**C**) Δ*ryhB* cells (JC1323) overexpressing CyaR, ChiX or McaS sRNA from the pBRplac plasmid, RNase T or YhbS from the pQE-80L plasmid, or AspX or YicC from the pBR* plasmid. Cells were induced with 50 μM IPTG at inoculation for 6 h at 37°C; mCherry fluorescence was measured at the endpoint. Biological triplicates were measured for each sample, and data are presented as mean and standard deviation. Whole cell lysates and total RNAs were prepared from cultures in **Fig. 3B** at the endpoint and analyzed by Western blot (**D**) and Northern blot (**E**), respectively. Relative quantitation of mCherry protein and RyhB sRNA is presented in **Fig. S3G** and **S3H** respectively. Unpaired two-tailed Student’s t-test was used to calculate statistical significance. Not significant (ns), P > 0.05; (*) P < 0.05; (**) P < 0.01; (***) P < 0.001.

In addition to measuring fluorescence signals, we also measured mCherry protein levels in cells carrying the plasmids (**Fig. 3D**). In line with fluorescence readings, overproduction of each factor increased mCherry protein levels by 2.5-to 9.4-fold (**Fig. S3G**). We further measured the effect of the plasmids on endogenous *sodB* expression. We engineered a FLAG tag to the C-terminus of SodB, and measured SodB-FLAG protein as well as *sodB* mRNA in cells overexpressing individual factors (**Fig. 3D, E**). Overproduction of the individual factors greatly increased SodB-FLAG, and the stimulatory effects were generally parallel to that of SodB-mCherry, with the exception of YhbS. Overproduction of YhbS led to a clear increase in *sodB* mRNA (**Fig. 3E**), but only a modest increase in SodB-FLAG compared to SodB-mCherry (**Fig. 3D**). We do not currently understand this discrepancy. It might reflect the fact that the *sodB*-mCherry and *sodB*-FLAG genes are encoded at different genomic loci and driven by different promoters. It is also possible that YhbS overproduction has pleiotropic effects on *sodB* expression at its native locus, such as inhibiting its translation elongation, in addition to antagonizing RyhB repression. Consistent with the observed increase in SodB protein, overproduction of each factor also increased *sodB* mRNA levels, though to a lesser extent (**Fig. 3E**). Collectively, these data demonstrate that the seven factors are *bona fide* antagonists for RyhB-mediated *sodB* repression in *E. coli*.

We also measured RyhB sRNA accumulation in these cells. RyhB levels were significantly lower in cells expressing individual factors compared to that of vector controls, with YicC overproduction resulting in the most dramatic decrease (**Fig. 3E**, quantification shown in **Fig. S3H**). As RyhB is constitutively expressed in *fur-*cells, it seems unlikely that the factors affect the synthesis of the sRNA. This was directly tested using a transcriptional fusion of the *ryhB* promoter to *lacZ*, in a Δ*fur* strain (**Fig. S3I)**. Only YhbS had a modest effect on this fusion, reducing it by half.

If the factors interfere with RyhB regulation, either by decreasing RyhB levels or blocking its function, then other RyhB targets beyond *sodB* should be affected as well. RyhB regulates more than a dozen mRNAs, including *sdhC* and *fumA* (19, 26). We constructed fluorescent reporters for *sdhC* and *fumA*, and measured fluorescence signals in cells expressing each of the seven factors. Notably, the *sdhC* reporter fusion had drastically reduced fluorescence levels in the *fur* deletion, similar to what was seen for *sodB* (**Fig. S4A**). Overexpression of individual factors in the *sdhC* reporter strain led to effects similar what was seen in the *sodB* reporter (**Fig. S4B**). Northern blot analysis of endogenous *sdh* polycistronic mRNA (*sdhCDAB-sucABCD*) also showed similar effects to those for the *sodB* mRNA; overproduction of ChiX or YhbS strongly increased *sdh* mRNA levels (**Fig. S4C**). The *fumA*-mCherry reporter showed less dramatic changes in a *fur* null strain when individual factors were overexpressed (**Fig. S4D**). CyaR, ChiX, McaS or RNase T showed modest increases, and YhbS slightly decreased fusion expression. AspX or YicC greatly increased fluorescence, comparable to what was seen for the *sodB* reporter. Taken together, these results suggest that each genetic factor, when overproduced, can suppress RyhB regulation of multiple mRNA targets, and the regulatory efficacies are not directly correlated with RyhB levels (for *yicC, aspX*, and *rnt*, for instance). We assume here that changing the specific reporter did not change the levels of RyhB.

### Endogenous genes modestly affect RyhB-mediated *sodB* expression

We then examined effects from the chromosomal copy of each of the individual regulators on RyhB signaling, by comparing *sodB*-mCherry expression in WT versus cells with single deletions of individual factors (**Fig. S5A**). The same deletions were also tested in the absence of RyhB (**Fig. S5B**). Cells deleted for *chiX, cyaR* or *yicC* modestly decreased SodB-mCherry production; these effects were dependent upon RyhB (compare **Fig. S5A** and **S5B**). Deletions of *rnt* or *aspX*, however, increased mCherry fluorescence either in the presence or absence of RyhB (**Fig. S5A, B**). Notably, these deletions also significantly slowed cell growth, which likely contributed to the apparent increase in signal. The *aspX* gene overlaps with the *aspA* coding sequence; thus its deletion affected AspA protein production as well. The role of chromosomally encoded AspX is further discussed below. Collectively, these data suggest that chromosomal copies of some of these genes do contribute to RyhB-dependent regulation, and likely would have stronger effects under the appropriate growth conditions. Deletion of *mcaS* or *yhbS* had no significant effects under the tested growth condition.

### General and RyhB-specific functions of individual regulators

These regulators were identified by studying their effects on RyhB function. We further determined if the effects from overproduction of individual regulators were specific to RyhB or affected other sRNA and Hfq-dependent regulation as well. We tested these regulators using the *chiP*-mCherry fusion (**Fig. 4A**), the *rpoS*-mCherry fusion (**Fig. 4B**), the *mutS*-mCherry fusion (**Fig. S6A**) and a control RBS1-mCherry fusion (**Fig. S6B**). In addition, we directly measured the levels of the sRNAs CyaR and ArcZ in cells overexpressing individual regulators (**Fig. 4C**). The results of these assays, discussed below in the context of each factor, demonstrated that five of the factors affected regulation and sRNA levels beyond RyhB. Two of the factors, AspX and YicC, exhibited specificity to RyhB and its targets.

**Figure 4.**
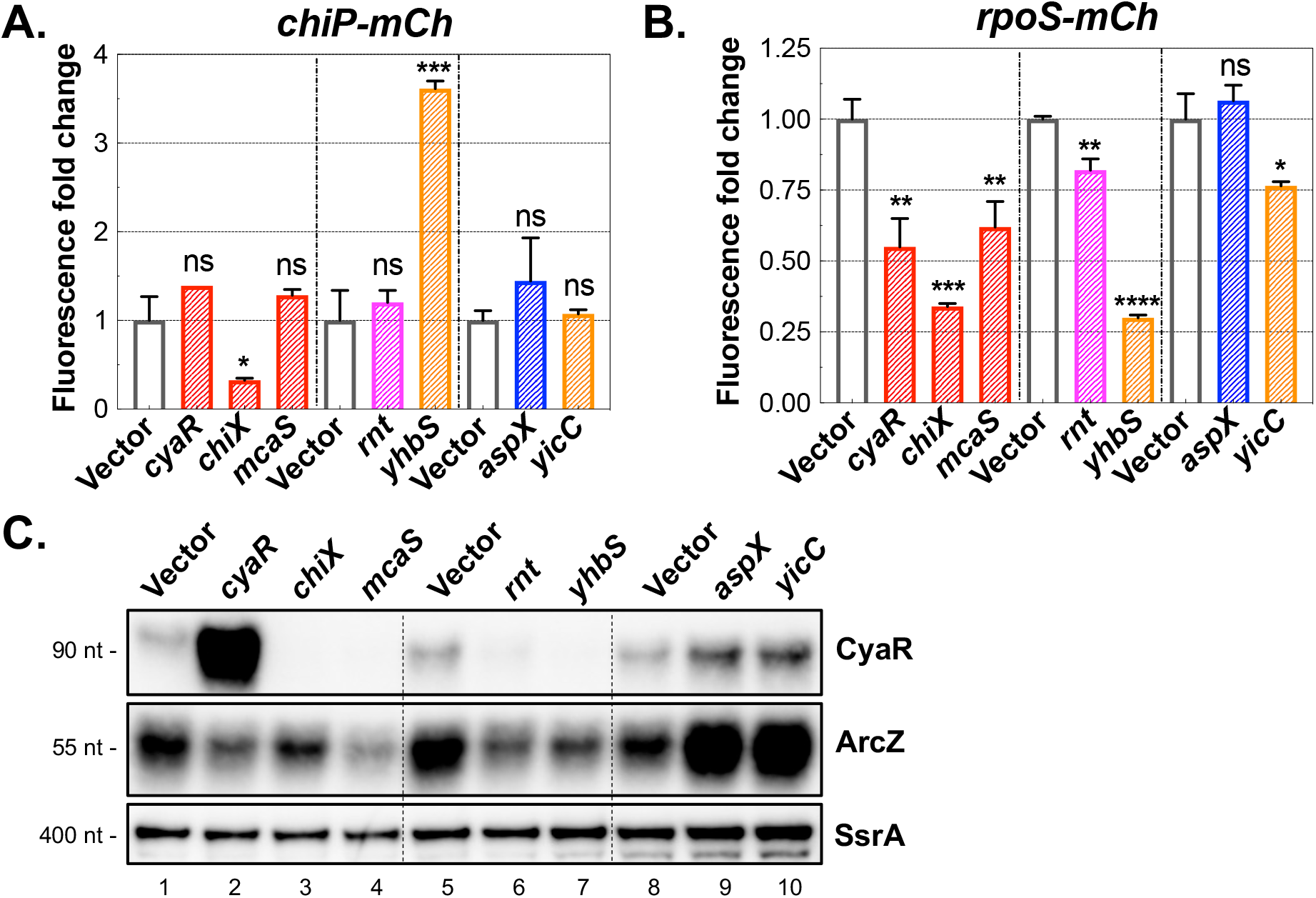
Many factors, especially ChiX and YhbS, affect stability and function of sRNAs beyond RyhB. Individual regulators, including CyaR, ChiX or McaS sRNA expressed from pBRplac, or His-RNase T or His-YhbS from pQE-80L, or AspX or YicC from pBR* were transformed into the reporter strains (**A**) *chiP*-mCherry (JC1247) or (**B**) *rpoS*-mCherry (JC1329) and grown for 6 h at 37°C with inducer (50 μM IPTG). mCherry fluorescence was measured at the endpoint using a TECAN Spark 10M microplate reader. The *chiP* reporter is repressed by ChiX and *rpoS* translation is activated by three sRNAs (DsrA, RprA and ArcZ); all are expressed endogenously in these strains. **C**. Northern blots were used to measure sRNA levels for CyaR or ArcZ from RNA samples from **Fig. 3B**; SsrA served as an RNA loading control. The same RNA samples, probing RyhB and *sodB*, were shown in **Fig. 3E**. Relative quantitation of CyaR and ArcZ sRNAs are presented in **Fig. S6C**. Unpaired two-tailed Student’s t-test was used to calculate statistical significance. Not significant (ns), P > 0.05; (*) P < 0.05; (**) P < 0.01; (***) P < 0.001; (****) P < 0.0001.

### Hfq-dependent sRNAs ChiX, McaS and CyaR act as general regulators via competition for Hfq

Three factors, ChiX, McaS, and CyaR, are Hfq-dependent sRNAs; they belong to a subgroup of sRNAs (Class II) that bind Hfq on its proximal and distal surfaces (27). Some sRNAs, exemplified by ChiX, have previously been shown to affect stability and function of other sRNAs via Hfq competition (28-30), likely by affecting the availability of Hfq for binding to other sRNAs (29, 31, 32). If Hfq is limiting, one would expect other targets of sRNA-dependent regulation to be affected as well, and levels of other sRNAs to be decreased. Consistent with this, all three of these sRNAs decreased the ability of endogenous sRNAs to activate the *rpoS*-mCherry fusion (**Fig. 4B**), and decreased levels of sRNAs CyaR and ArcZ (**Fig. 4C**). Only ChiX released repression of the rather sRNA-independent *mutS*-mCherry fusion and none had an effect on the control RBS1 fusion (**Figs. S6A, B**).

Not all sRNAs have this effect. Overproduction of ArcZ, a Class I sRNA, or MgrR, another Class II sRNA, had no effect on *sodB*-mCherry fluorescence (**Fig. S7A**). Consistent with the idea that Hfq becomes limiting upon sRNA overexpression, an increase in the level of Hfq at least partially restored RyhB regulation in cells overexpressing CyaR or McaS (**Fig. S7B**). Hfq levels in these cells were increased by ∼2-fold (**Fig. S7C**), as seen previously (29). The Hfq plasmid improved RyhB regulation in cells overexpressing ChiX very modestly (**Fig. S7B**). However, the level of Hfq was high when ChiX was overexpressed even without the Hfq plasmid (**Fig. S7C**, compare lane 5 to lane 1 and 3), and did not significantly increase with the Hfq plasmid. This suggests that ChiX overexpression abrogates the ability of Hfq to negatively regulate its own expression. This autoregulation is known to be dependent on the distal face of Hfq (33). Interestingly, these results suggest that ChiX is able to interfere with RyhB regulation of *sodB*, positive regulation of *rpoS* and regulation of *mutS* (**Fig. 4, Fig. S6**), even when Hfq levels are high.

Overall, our data support that overproduction of these three Hfq-dependent sRNAs, ChiX, McaS and CyaR, can efficiently titrate Hfq from RyhB and likely the *sodB* target mRNA as well, leading to a defect in regulation by RyhB. We noted that although not dramatic, deletion of either *chiX* or *cyaR* was sufficient to improve *sodB*-mCherry repression, dependent upon RyhB (**Fig. S5**). We speculate that under conditions that significantly induce these competitor sRNAs, these sRNAs may expand their regulons through Hfq competition.

### Exonuclease RNase T overproduction has a general effect on sRNA-dependent regulation

In our original *sodB* fusion screen, we identified three distinct *rnt* clones that increased reporter fluorescence production (**Figs. 2 and S3E**). Overexpression of RNase T led to potent suppression of RyhB function and lower sRNA levels (**Figs. 3 and 4**). As for the Class II sRNAs discussed above, overexpression of RNase T modestly disrupted *rpoS*-mCherry activation (**Fig. 4B**), led to a significant decrease in levels of CyaR (**Figs. 4C and S6C**), but did not disrupt *mutS* regulation (**Fig. S6A**). Its behavior suggests that RNase T is a general regulator of sRNAs, albeit not as effective as the best of the sRNAs, ChiX.

RNase T, a 3′-5′ exoribonuclease, is responsible for the 3′end processing of many stable RNAs, including tRNAs. It can trim stable RNA substrates to duplexes with a short 3′-overhang of 1 to 4 nts (34). RNase T mutants defective in either substrate binding (R13A) or catalysis (D23A) (35) lost the ability to relieve RyhB-dependent repression of *sodB*-mCherry (**Fig. 5A**). All RNase T variants were expressed at similar levels (**Fig. S8A**). These results demonstrate that the nuclease activity of RNase T is required for antagonization of RyhB regulation in cells.

**Figure 5.**
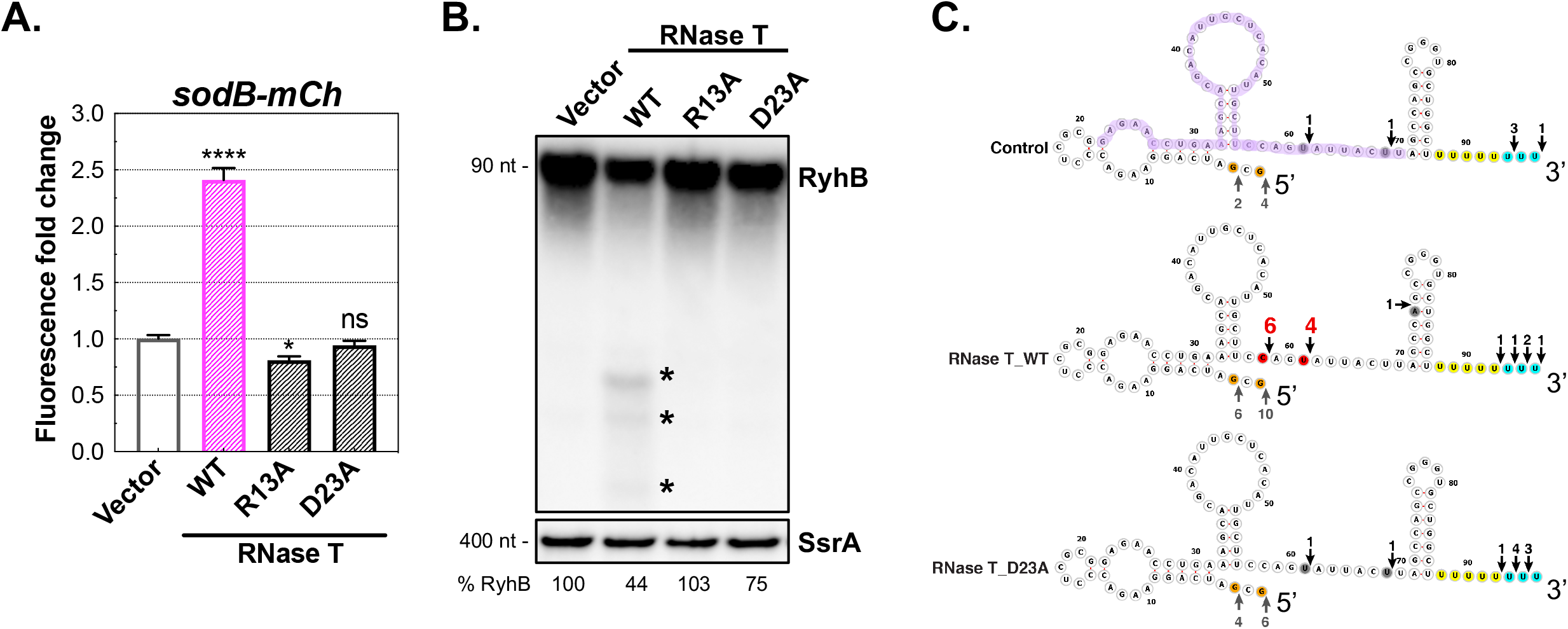
High levels of RNase T suppress RyhB sRNA function by promoting sRNA decay. **A**. Plasmid-borne WT or RNase T mutants (R13A: substrate-binding deficient; D23A: non-catalytic) in the *sodB* reporter strain (JC1249) were induced with IPTG (50 μM) for 6 h prior to fluorescence measurement. The RNase T constructs all carry an N-terminal His tag. **B**. *sodB* reporter strain (JC1249) transformed with WT or RNase T mutants were grown to OD_600_ of ∼2.0 in LB+Amp+IPTG (50 μM), and total RNAs were extracted for Northern blot analysis. RyhB fragments detected in lane 2 are highlighted with asterisks. Relative levels of the full-length RyhB are shown, normalized to the SsrA loading control first and then further compared to that of the vector control. Unpaired two-tailed Student’s t-test was used to calculate statistical significance. Not significant (ns), P > 0.05; (*) P < 0.05; (****) P < 0.0001. **C**. Total RNAs from cells expressing vector, WT or D23A of RNase T in **Fig. 5B** were subjected to 5′ and 3′ end analysis of RyhB transcripts, by RNA self-ligation, reverse transcription, cloning and sequencing as described in Materials and Methods. RyhB 5′ and 3′ ends are indicated by upward and downward arrows respectively. Number of clones mapped to a specific nucleotide position are labeled above/below each arrow, with the significant alterations highlighted in red. RyhB structure shown is based on (65). Purple highlight shows region detected by the probe used in **Fig. 5B**. Six or more 3′ U residues have been shown to be important for optimal Hfq binding (66) and are highlighted here in yellow; additional U residues found in RyhB are highlighted in blue.

Notably, RyhB levels were reduced to about 40% of the vector control, and shorter RyhB fragments (asterisks, **Fig. 5B**) were detected in cells overproducing the functional RNase T, likely representing degradation intermediates. We mapped the 5′ and 3′ ends of RyhB transcripts in cells with and without RNase T overproduction, using a method involving 5′-3′ self-ligation of RNA, reverse transcription, PCR cloning and sequencing (**Fig. S8B**) (36). In control cells expressing the vector plasmid or a mutant RNase T (D23A), the majority of the detected RyhB transcripts were full-length RNAs (4/6 in vector control and 8/10 in D23A) (**Fig. 5C**). In contrast, only 5/16 were detected as full-length RNAs in cells overproducing RNase T_WT, consistent with the decrease in full length RyhB detected in Fig. 5B. The other 11/16 RyhB transcripts were the 5′ halves of RyhB, presumably representing RyhB degradation intermediates. The truncated RyhB is likely defective for regulation of target genes and may be rapidly further degraded. As several other sRNAs also had reduced levels when RNase T was overexpressed (**Fig. 4C**), RNase T may contribute to the homeostasis of many sRNAs in cells.

RNase T exonuclease is blocked by hairpin structures but can cleave near such double-stranded structures (34, 37). Therefore, we expected to see shortening of the polyU tail of the RyhB terminator, rather than the observed 5′ halves of RyhB (**Fig. 5**). In fact, high levels of RNase T led to accumulation of modestly shorter RyhB transcripts in the absence of Hfq (**Fig. S8C**). Based on these observations, we favor a model in which higher levels of RNase T trim the polyU tail of the sRNA, compromising optimal Hfq binding, and subsequently allowing sRNA decay by other ribonucleases (**Fig. 8**). The *rnt* mutant has a significant growth defect in the tested condition, making it difficult to assess the contribution of the endogenous RNase T to sRNA stability and function.

### An uncharacterized GNAT family acetyltransferase acts as a general regulator of Hfq-dependent regulation

YhbS, a putative GNAT family acetyltransferase, was identified as a silencer of RyhB signaling in our genetic screen (**Figs. 2 and S3E**). Intriguingly, YhbS overproduction decreased levels of all sRNAs tested, including CyaR and ArcZ (**Fig. 4C**). It also disrupted functions of Hfq-dependent sRNAs, including ChiX-mediated *chiP* repression (**Fig. 4A**) and sRNA-mediated *rpoS* activation (**Fig. 4B**), as well as sRNA-independent repression of *mutS* by Hfq (**Fig. S6A**) and autoinhibition (**Fig. S9**). These results suggest an effect of YhbS on Hfq itself rather than on sRNAs. Consistent with YhbS targeting Hfq, a recent study reported YhbS association with Hfq in Hfq pulldown experiments (38). There are 26 acetyltransferases in *E. coli*, and many of them are not characterized (39). Given the possible functional redundancy, it is not surprising that deletion of *yhbS* alone was not sufficient to disrupt RyhB regulation (**Fig. S5**). Nevertheless, the biochemical and genetic interactions support a role of YhbS in acetylating Hfq or proteins associated with Hfq, which in turn perturbs Hfq/sRNA signaling (**Fig. 8**). Future work will explore this interesting protein.

### AspX, a small RNA expressed from the 3′end of *aspA*, specifically antagonizes RyhB function via an ‘RNA sponge’ mechanism

Our genetic screen identified multiple positive clones consisting of the intergenic region between the 3′ end of *aspA* and the 5′ end of *dcuA*, as well as portions of both genes (*dcuA*’<*Ig*<*aspA*’) (**Fig. S3E**). RNA-Seq data from the GWIPS-viz Genome Browser database (40) strongly suggest the presence of an unannotated short transcript encoded at the 3′end of *aspA* (red dashed box in **Fig. S10A**). Differential RNA-Seq data of the *E. coli* transcriptome (41) further suggests at least two primary and one processed transcript encoded at this locus (**Fig. S10B**). We named these transcripts AspX. Overexpression of AspX did not affect *rpoS*-mCherry activation (**Fig. 4B**), *chiP*-mCherry or *mutS*-mCherry repression (**Figs. 4A** and **S6**), and rather than decreasing levels of other sRNAs, as with the general factors discussed above, led to accumulation of CyaR and ArcZ (**Figs. 4C** and **S6C**). This increase may result from reduction in RyhB levels, freeing more Hfq to engage and stabilize other sRNAs. Therefore, AspX appears to be a RyhB-specific regulator.

AspX 5′ ends were mapped by 5′ Rapid Amplification of cDNA Ends (5′RACE); results are summarized in **Fig. S10C**. 5′RACE mapped a predominant transcription start site ‘A’ (Green color in TAP+ sample, corresponding to the P2 site in **Fig. S10B**), and a processed start site ‘C’ (Blue in TAP-sample). Based on the RNA seq signals and the intrinsic terminator sequence, these transcripts were predicted to be ∼118 bp (<u>s</u>hort, AspXS) and ∼98 bp (<u>p</u>rocessed, AspXP) respectively. A longer primary transcript AspXL (long, ∼278 bp), which has been detected by dRNA-Seq but not by 5′ RACE, was also included. Northern blots detected two major transcripts corresponding to the sizes of AspXL and AspXS from the chromosomal locus in WT cells (**Fig. S10D**, left panel, lane 1). In the absence of RyhB, transcripts corresponding to the sizes of AspXL and AspXP were seen, suggesting that RyhB may protect AspXS from processing (**Fig. S10D**, left panel, compare lanes 1 and 2). However, deletion of *aspX* showed no effect on endogenous RyhB levels, either in the absence or presence of Fur (Fig. S10D). Deletion of *hfq* in *fur* ^-^ cells greatly decreased RyhB levels (**Fig. S10D**, left panel, compare lane 3 and 5) but had little effect on AspX accumulation, suggesting AspX, unlike most sRNAs, is not stabilized by Hfq. Consistent with this, AspX was also not significantly enriched in Hfq immunoprecipitants (42). A summary of the genomic context of *aspX* is presented in **Fig**. S10E.

To characterize AspX regulation of RyhB, we expressed the three AspX isoforms in the *sodB* reporter strain and measured RyhB levels as well as *sodB*-mCherry fluorescence (**Figs. S11A, B**). Overexpression of AspXL or AspXS reduced RyhB levels and increased *sodB*-mCherry fluorescence, whereas AspXP showed no effect. Interestingly, RIL-Seq data, in which the direct interactions of sRNA-mRNA partners on Hfq can be captured, suggested abundant interactions between RyhB and the *aspA* 3′end (now named AspX) under an iron starvation condition (42). This result hinted that AspX might control RyhB levels and likely functions through direct base-pairing interactions. The Mfold web server (43) predicted a perfect 10-nt stretch of pairing (**Fig. 6A**) between sequences at the terminator loop and flanking region of RyhB (nt 73-82, cyan in **Fig. 6B**) and a region near the 5′ end of AspXS (nt 13-22). Notably, the putative RyhB region for pairing with AspXS is different from the RyhB primary seed sequence for pairing with known mRNA targets (nt 37-51, yellow in **Fig. 6B**). Interestingly, AspXP is missing the predicted pairing sequence, likely due to an endonucleolytic cleavage event (a red triangle in **Fig. S10E** indicates the cleavage site), and this may explain the loss of suppression of RyhB regulation by the AspXP RNA (**Fig. S11B**). We confirmed the direct base pairing interaction between AspX and RyhB using two mutants in AspXL (m_GCGC_ and m_C_) that should disrupt the predicted pairing and two RyhB compensatory chromosomal mutants (*ryhB*_gcgc and *ryhB*_g; **Fig. 6A**). These were tested for *sodB*-mCherry fluorescence as well as cellular RyhB levels (**Fig. 6C and D**). Overproduction of AspX in *ryhB*_WT cells drastically reduced RyhB levels and increased *sodB*-mCherry fluorescence. Base-pairing mutants of AspX or mutations in RyhB alone abrogated downregulation of RyhB and upregulation of *sodB*-mCherry. Only the AspX mutants that restore pairing with RyhB mutants were able to decrease RyhB levels and significantly increase *sodB*-mCherry fluorescence (**Figs. 6C, 6D**, lanes 7 and 12). Therefore, AspX uses a short sequence to base pair with the terminal stem-loop of RyhB, leading to loss of RyhB and thus its regulatory function.

**Figure 6.**
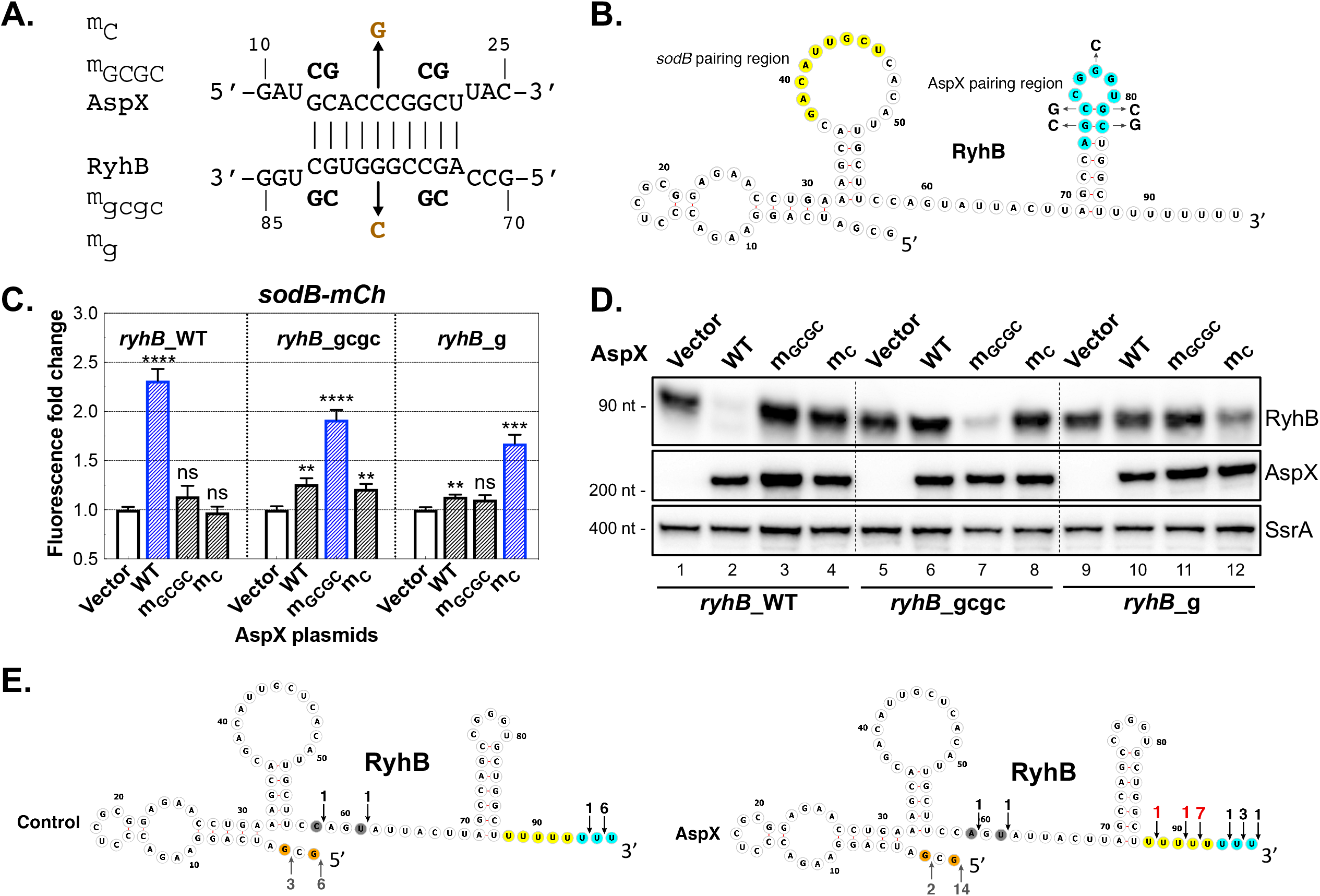
AspX, a short RNA expressed from the 3′end of *aspA* gene, selectively antagonizes RyhB function via an ‘RNA sponge’ mechanism. **A**. Predicted base pairing of RyhB and AspXS. Nucleotide changes of two AspX mutants and corresponding compensatory RyhB chromosomal mutations are shown. **B**. Secondary structure of RyhB sRNA (65), with the pairing sequences highlighted in yellow (the primary seed region for mRNA pairing (65)) and in cyan (the predicted region to pair with AspX). **C**. RyhB chromosomal variants (WT: JC1291 or mutant *ryhB*_gcgc: JC1292 or *ryhB*_g: JC1293) were transformed with WT or mutant AspXL and induced with IPTG (50 μM) for 15 h at 30°C before measuring mCherry fluorescence. Unpaired two-tailed Student’s t-test was used to calculate statistical significance. Not significant (ns), P > 0.05; (**) P < 0.01; (***) P < 0.001; (****) P < 0.0001. **D**. Total RNAs were prepared from cultures in **Fig. 6C**, and analyzed by Northern blot for RyhB, AspX and SsrA. **E**. Total RNAs of control or AspX from **Fig. 6C** were used to map RyhB 5′ and 3′ ends as for **Fig. 5C**. RyhB 5′ and 3′ ends are indicated by upward and downward arrows respectively. Number of clones mapped to a specific nucleotide position are labeled above/below each arrow, with the significant alterations highlighted in red. Six or more 3′ U residues have been shown to be important for optimal Hfq binding (66) and are highlighted here in yellow; additional U residues found in RyhB are highlighted in blue.

The identification of these pairing mutations in AspX allowed us to create a chromosomal *aspX* mutant, *aspX*_GCGC_, that did not have the growth defect previously seen for Δ*aspX*. However, this chromosomal mutant had no measurable effect on the expression of the *sodB*-mCherry fusion, either in a *fu*r^+^ or *fur*^-^ strain (**Fig. S10F**)

AspX pairing led to an overall decrease of RyhB levels as well as accumulation of slightly shorter transcripts (**Figs. 3E, 6D and S11C**). RNA samples from **Fig. 6D** were used to determine the 5′ and 3′ ends of the RyhB transcripts, using the methodology described earlier (**Fig. S8B**). The majority (9/16) of RyhB in AspX-overproducing cells had intact 5′ ends but shortened 3′ polyU tails (n ≤ 5) relative to the control sample (**Fig. 6E**), suggesting that AspX regulates the polyU tail length of RyhB.

Hfq was reported to participate in RNA sponge-mediated regulation of sRNAs (8, 44). In a strain deleted for *hfq*, RyhB levels were low and AspX overproduction did not further decrease them or lead to a shorter RNA (**Fig. S11C**), suggesting that Hfq is required for AspX regulation of RyhB. AspX exhibits anti-RyhB activity via a base-pairing mechanism; thus it is a new RNA sponge or decoy. Only RyhB was found to form an RNA duplex with AspX on Hfq (42), supporting that AspX is a RyhB-specific RNA sponge.

Other RNA sponges for RyhB have been reported, likely reflecting the importance of RyhB and RyhB-like sRNAs for iron homeostasis. Lalaouna *et al* (10) reported that a byproduct of a tRNA precursor, named 3′ETS^*leuZ*^, acts as a sponge to reduce RyhB transcriptional noise in *E. coli*. Han *et al* (9) described *skatA*, a fragment from the 3′end of *katA* mRNA of *Pseudomonas aeruginosa*, that base pairs with PrrF1, a RyhB functional homolog, and disrupts its ability to regulate target mRNAs. RNA sponges have been reported for other sRNAs as well, including ChiX (18, 45), GcvB (44), FnrS (11) and RybB (8, 10). Emerging evidence from RNA proximity ligation-based high throughput sequencing analyses, RIL-Seq and GRIL-Seq (8, 9), have suggested that RNA sponge-mediated modulation of sRNA activities is a widespread mode of bacterial gene regulation. Our fluorescence-based genomic library approach provides an effective way to identify RNA sponges, as illustrated by identification of AspX as a new RNA sponge for RyhB, and the *chbBC* IGR decoy for ChiX.

While this work clearly defines the ability of AspX to silence RyhB, a number of interesting questions remain. AspX is transcribed by promoters inside the *aspA* coding sequence (**Fig. S10**), and the regulation of those promoters and thus the conditions when AspX may be most important are not yet known. Once made, the short AspX transcript is relatively stable. The detection of AspX-RyhB chimeras via RIL-Seq under iron starvation conditions (42) and the endogenous RyhB-dependent protection of AspXS from cleavage to the inactive AspXP (**Fig. S10D**) suggest that AspX is available and interacts with RyhB under these growth conditions. A putative RNase E cleavage is likely to generate the shortest isoform AspXP (46), and RyhB seems to block RNase E-mediated AspX cleavage via base pairing at the cleavage site (**Fig. S10**), thus forming a self-sustained and self-contained regulatory loop for RyhB and AspX.

### YicC, a conserved protein, selectively leads to rapid RyhB degradation through PNPase

Gene deletion and overexpression analyses demonstrated that the hypothetical protein YicC was necessary and sufficient to disrupt RyhB signaling, leading to *sodB* derepression (**Fig. 3**). Of the seven regulators, YicC overproduction led to the greatest reduction of RyhB (**Fig. 3E**) but had little effect on other tested sRNAs or their targets (**Figs. 4 and S6**). Therefore, YicC acts like a specific RyhB regulator.

YicC homologs are widely present in both Gram-negative and positive bacteria (**Fig. S12A**), defined by a highly conserved N-terminal YicC-like domain and a C-terminal DUF1732 domain. Little is known about their function. In *E*.*coli*, YicC was one of many genes implicated in stress-induced mutagenesis (47) and cells carrying insertions in both *yicC* and the neighboring *dinD* gene showed defective growth in stationary phase or at high temperatures (48). In a time course experiment, YicC induction resulted in a rapid decrease of RyhB levels (**Fig. 7A**). As YicC expression had little effect on *ryhB* transcription (**Fig. S3I**), this rapid loss of RyhB suggested that YicC might activate an RNA decay pathway for RyhB degradation.

**Figure 7.**
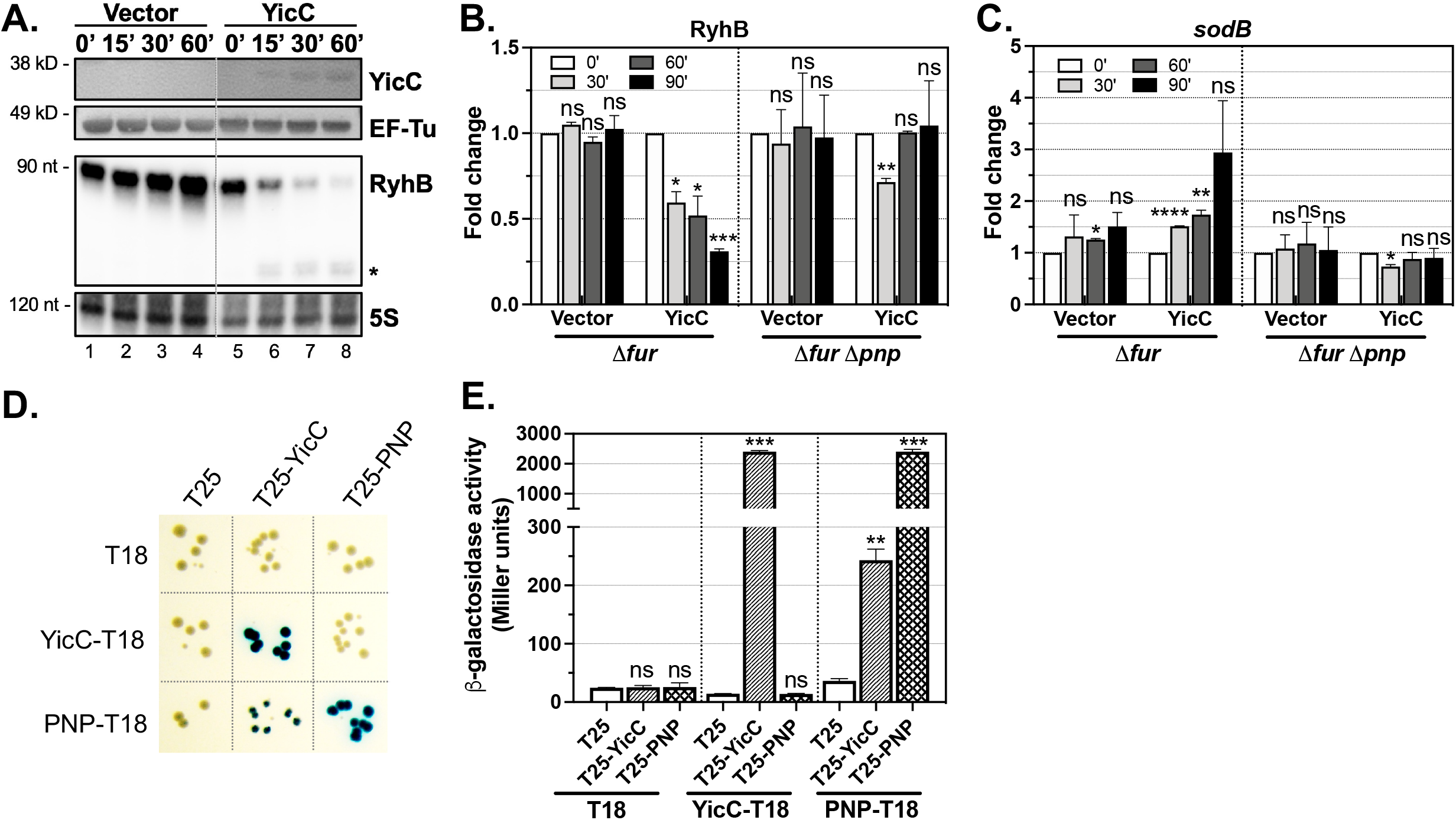
YicC collaborates with PNPase to selectively degrade RyhB. **A**. The *sodB* fluorescence fusion strain (JC1249) harboring empty vector (pQE-80L) or pYicC was grown to OD_600_ of ∼0.15 in LB+Amp, and then induced with 100 μM IPTG for 0, 15′, 30’, 60’. Total RNA and protein samples were prepared from cultures at the indicated timepoints and analyzed by Western and Northern blotting. EF-Tu served as a protein loading control and 5S rRNA served as an RNA loading control. **B**. WT (XL111) or Δ*pnp* (XL97) transformed with empty vector (pBR*) or pYicC2 were grown to OD_600_ ∼0.3 in MOPS Neidhardt EZ rich medium, IPTG was added to a final concentration of 200 μM, and sample cultures were taken at 0’, 30’, 60’ and 90’ for RNA and protein extraction. Levels of YicC induction are shown in **Fig. S12D**. Total RNA samples were analyzed for RyhB, MicA, ArcZ, GcvB, *sodB* and SsrA by Northern blotting (shown in **Fig. S12E**). Quantification of the Northern blot for RyhB is in **Fig. 7B** (biological duplicates) and for *sodB* in **Fig. 7C**. Quantitation for other sRNAs by Northern blot is in **Fig. S12F**. Data are mean and standard deviation. **D**. BACTH reporter strain BTH101 co-transformed with plasmids of T18 and T25 fused to YicC or PNPase was spotted on LB agar supplemented with Amp, Kan and X-gal, and bacterial colonies were imaged after incubation at 30°C for 60 h. **E**. BACTH transformants were grown for 16 h at 30°C in LB supplemented with Amp and Kan, and assayed for β-galactosidase activity. Biological triplicates were assayed, and data are presented as mean and standard deviation. Unpaired two-tailed Student’s t-test was used to calculate statistical significance. Not significant (ns), P > 0.05; (*) P < 0.05; (**) P < 0.01; (***) P < 0.001; (****) P < 0.0001.

**Figure 8.**
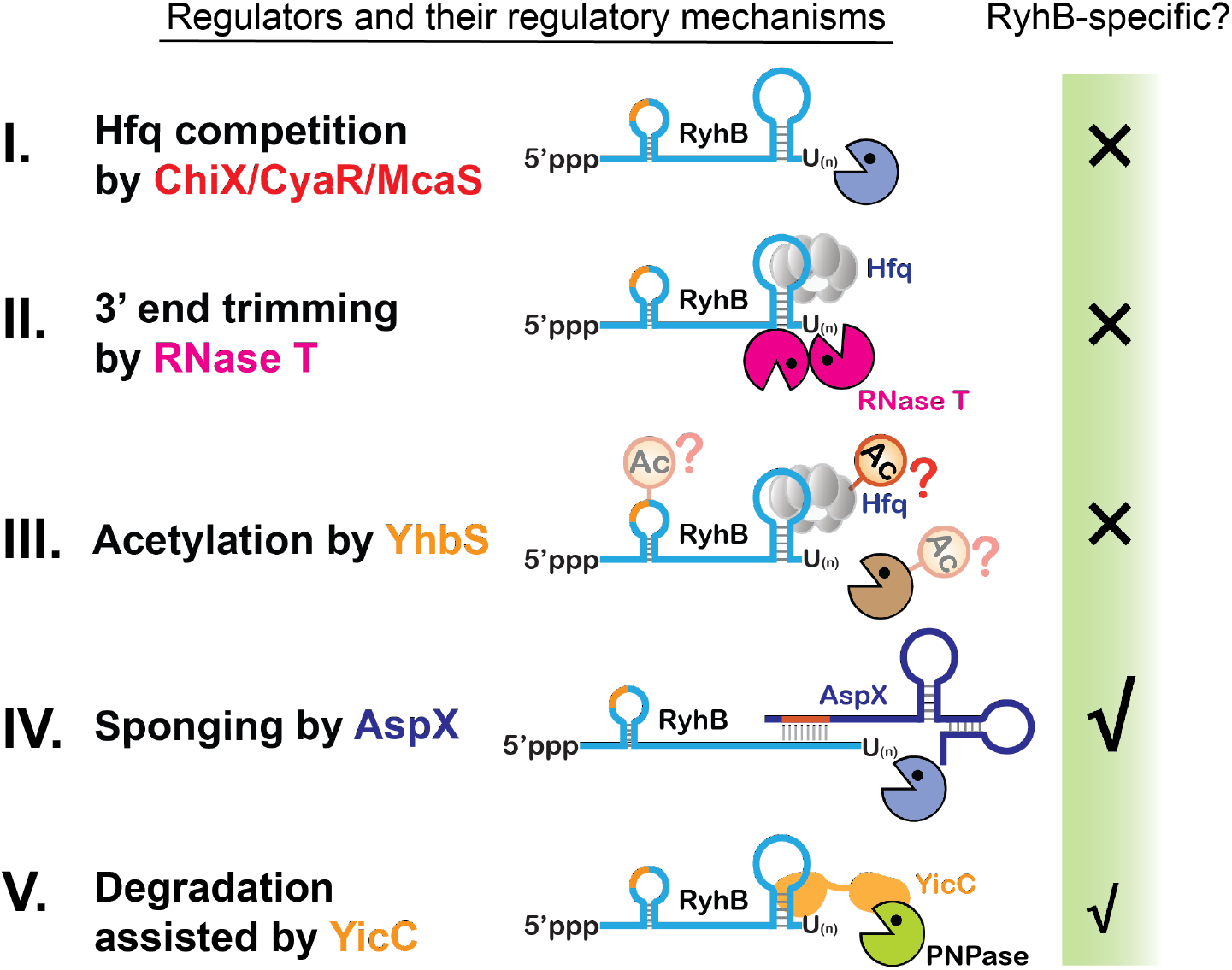
Diverse regulators silence sRNA signaling in *E. coli*. **I**. Three class II sRNAs ChiX, CyaR and McaS sequester the RNA chaperone Hfq, leading to decay of sRNAs that are not bound to Hfq, including RyhB. **II**. Exonuclease RNase T competitively trims sRNA polyU tails, thus compromising Hfq binding, leading to sRNA degradation. **III**. YhbS, a putative acetylase, acts broadly on sRNA/Hfq signaling, possibly by acetylating Hfq or proteins in the Hfq complexes. **IV**. AspX forms complementary base pairing with the RyhB 3′ terminator to destabilize RyhB; AspX appears to selectively repress RyhB function. **V**. YicC, a stress-induced protein, selectively degrades RyhB and likely a few other sRNAs via the 3′-5′ exoribonuclease PNPase.

Endoribonucleases RNase E and RNase III have been reported to cleave RyhB sRNA and its mRNA targets (49-52). We tested if YicC could silence RyhB function via either ribonuclease, comparing the ability of YicC to disrupt RyhB regulation in WT versus ribonuclease deficient cells (**Fig. S12B**). In WT strains, overproduction of YicC increased *sodB*-mCherry fluorescence by 3.3-fold; in the RNase E mutant lacking the C-terminal degradosome platform (*rne131*) or the RNase III mutant (*rnc14*), *sodB*-mCherry fluorescence in the vector control was slightly increased. Overexpression of YicC still led to significant increases of reporter fluorescence, suggesting neither nuclease is essential for YicC-mediated disruption of RyhB regulation. Given that RNase E is essential in *E. coli*, we cannot rule out that the N-terminal portion of RNase E might be involved. Consistent with the reporter assay, RyhB disappeared when YicC was overproduced in *rnc14* cells, as it did in WT (**Fig. S12C**). Notably, ChiX sRNA levels were not affected by YicC overproduction, consistent with our finding that YicC did not disrupt ChiX-dependent repression (**Fig. 4A**).

The 3′-5′ exoribonuclease PNPase also participates in RyhB sRNA metabolism (53, 54). Thus, we compared RyhB accumulation and activity in control and YicC overproduction cells in a WT or *pnp* null background. The *pnp* null mutant grew poorly in LB medium. To improve bacterial growth as well as plasmid retention in the *pnp* mutant, we expressed YicC from a lower copy number plasmid in EZ rich defined medium; YicC proteins were expressed at similar levels in the WT and *pnp* mutant in these conditions (**Fig. S12D**). With this plasmid, YicC induction in WT cells led to a dose-dependent decrease of RyhB (see Northern blot in **Fig. S12E** and quantitation in **Fig. 7B**). Strikingly, in the *pnp* null cells, there was no decrease in RyhB levels after YicC induction. At the same time, *sodB* levels, which increased upon YicC expression in the WT strain, did not increase in the *pnp* mutant (**Figs. 7C** and **S12E**), confirming a requirement for PNPase for YicC function.

As noted above, YicC overproduction did not affect levels of ChiX (**Fig. S12C**) but upregulated CyaR or ArcZ (**Figs. 4C and**,**S9C**). After a shorter induction in the EZ rich medium, ArcZ and GcvB sRNAs were unaffected by YicC in either the WT or *pnp* mutant (**Fig. S12E, F**). However, YicC overproduction led to a reduction of MicA sRNA, again dependent on PNPase (**Fig. S12E, F**). Therefore, YicC shows significant selectivity, decreasing the accumulation of a subset of Hfq-dependent sRNAs.

The dependence of YicC on PNPase suggested that YicC may serve as a co-factor or adaptor, bringing specific sRNAs to PNPase. Thus, we examined possible YicC-PNPase interaction using a bacterial two hybrid (BACTH) assay in which each protein was fused to the T18 or T25 fragments of adenylate cyclase (**Figs. 7D, E**). We detected a significant interaction between PNP-T18 and T25-YicC, both by a qualitative test (**Fig. 7D**) and a quantitative assay (**Fig. 7E**). In the opposite orientation, no interaction was detected; it seems likely that the protein tags interfere. The BACTH assay also detected strong self-interactions of PNPase and YicC.

PNPase, found in both prokaryotes and eukaryotes, is known to associate with other proteins, changing its degradative capabilities. In *E. coli*, it associates with RNase E as a core component of the bacterial degradosome, working with other degradosome components, including RNA helicase RhlB; it also associates with RhlB in the absence of the degradosome (55, 56). In a number of bacteria, PNPase associates with Y RNA and the bacterial Ro protein, with these accessory factors modulating what is degraded by PNPase (57). Here, our data support that YicC, a highly conserved protein of previously unknown function, directs PNPase to selectively degrade RyhB. There are accumulating examples of ribonucleases that selectively degrade one or one group of sRNAs via collaboration with a cofactor. RNase E is reported to selectively degrade GlmZ or CsrB/C sRNA through collaboration with the adaptor protein RapZ (58) or the GGDEF-EAL domain protein CsrD (59) respectively. Given the wide conservation of YicC, it seems quite possible this association with PNPase may have conserved functions in many bacteria.

While our study identified YicC function when it is overproduced, deletion of *yicC* had a modest but statistically significant effect on *sodB*-mCherry expression, dependent upon RyhB (**Fig. S5**), suggesting that this protein’s effects on RyhB are likely to be physiologically significant. Future work will focus on defining the conditions under which YicC is expressed, its *in vivo* roles and the molecular architecture of YicC-PNPase-RyhB interactions.

## Summary

sRNAs can act as potent modulators of stress responses in bacteria. When the stress is past or cells have adapted, bacteria then will turn down sRNA activities. In optimal growth conditions, cells also often keep sRNAs in check. Combining a genomic library approach with fluorescent reporters, we were able to readily identify both known and new regulators of sRNA signaling. Previously, De Lay and Gottesman (54) used a selection combined with a blue-white screen after general mutagenesis and successfully identified new sRNA regulators. El-Mowafi *et al*. (60) carried out a screen by using a plasmid-based fluorescent reporter and a library of in-cell produced cyclic peptides and successfully identified small peptide inhibitors of Hfq. These examples demonstrate that a robust reporter system in combination with a good screen strategy can serve as a very useful tool for discovery of new chemical or genetic modulators of sRNA signaling.

In this work, we identified a previously reported RNA sponge (18, 45) as a strong suppressor of ChiX signaling. In addition, we identified both previously known and new negative regulators of RyhB and RyhB activity. These regulators act via distinct regulatory mechanisms; some are specific to RyhB and others play broader roles in sRNA function. All regulators appeared to function posttranscriptionally, highlighting the complexity and versatility of controlling sRNA pathways beyond gene transcription. A single screen of one regulatory pair, RyhB repression of the *sodB* mRNA, yielded a novel RNA decoy and two uncharacterized protein regulators, suggesting that very likely many other regulators, particularly ones specific to other sRNAs, remain to be discovered.

## Materials and methods

### Bacterial strains and culture conditions

*E. coli* K-12 strain MG1655 and its derivatives were used and are listed in Supplemental Table S1. Plasmids are listed in Supplemental Table S2. Descriptions of strain and plasmid construction is in Supplementary Materials and Methods. DNA sequences, including PCR oligos, Northern probes and gBlocks gene fragments (Integrated DNA Technologies) are listed in Supplemental Table S3. Phage P1 *vir* transduction (61) was used to move genetic elements as needed. Ectopic expression of Hfq-dependent sRNAs and AspX variants were from the medium-copy number plasmid pBRplac (AmpR) and pBR* (KanR) respectively. Protein coding genes were typically expressed from the high-copy number plasmid pQE-80L (AmpR), with the exception of *yicC*, which was cloned to both pBR* and pQE-80L plasmids. The pQE-80L expressed proteins carry an N-terminal His tag.

Bacteria were grown in LB broth (Lennox), minimal M63 glucose medium (KD Medical, MD) or EZ rich defined medium (Teknova). Glucose was supplemented to a final concentration of 0.2% for M63 medium. Unless otherwise indicated, 0.1% L-arabinose or 50 μM IPTG (isopropyl β-D-1-thiogalactopyranoside) were used for induction of genes driven by the *pBAD* or *plac* promoter, respectively. Appropriate antibiotics were supplemented as needed for plasmid maintenance or for selection at the following concentrations: Ampicillin (Amp), 100 μg/ml; Kanamycin (Kan), 50 μg/ml; Zeocin (Zeo), 25 μg/ml; Chloramphenicol (Chl), 10 (for selection of *cat-sacB*) or 25 (for all others) μg/ml.

### Fluorescence-based genomic library screen

Parental reporter strains expressing basal fluorescence (*chiP* fusion: JC1200 and *sodB* fusion: JC1248) were transformed with 30 ng of the *E. coli* genomic plasmid library (62) via electroporation, and recovered cells were selected on M63 glucose agar supplemented with Amp. Appropriate dilutions were selected to provide ∼500 colonies per plate, for a total of 50 agar plates per screen. Colony fluorescence on agar was captured after growth at 37°C for 48 h, using the Bio-Rad ChemiDoc MP Imaging system (settings: Cy2 for GFP, Cy3 for mCherry and Ponceau S for Bright Field, auto optimal exposure). The majority of colonies had low basal fluorescence, and a few clones with visually brighter mCherry and/or GFP fluorescence were purified by re-streaking 2-4 times on selective agar until isolation of pure single clones.

Plasmids were then extracted and re-transformed into the parental reporter strain (JC1200 or JC1248) for verification of elevated fluorescence. Verified plasmids were sequenced from both ends using sequencing oligos pBRlib_For and pBRlib_Rev, and mapped to the *E. coli* K-12 genome using EcoCyc BLASTn search. In parallel, eight clones (C1-C8) that exhibited basal fluorescence were purified as negative controls. Qualitative measurement of colony fluorescence was conducted by using the ChemiDoc MP Imaging system and quantitative measurement of fluorescence was done in liquid culture using the TECAN Spark 10M microplate reader.

### Fluorescent reporter assays

To visualize colony fluorescence, *E. coli* cells were streaked on LB agar supplemented with appropriate antibiotics, and grown at 37 °C for 16 h. The next day, colonies on agar were directly imaged using the Bio-Rad ChemiDoc MP Imaging system at multi-mode (Cy2 for GFP, Cy3 for mCherry and Ponceau S for Bright Field, auto optimal exposure). Fluorescent images were processed using Image Lab 6.0 (Bio-Rad). To quantitate fluorescence signals, six biological replicates for each strain were inoculated into 96-well microplates (#3596, Corning, NY) with 100 μl LB supplemented with appropriate antibiotics and inducers in each well, grown at 37°C for 6 h with shaking (250 rpm), and then cell fluorescence as well as optical density (OD_600_) were measured using a TECAN Spark 10M microplate reader (GFP: 475 ± 20 nm ex.; 520 ± 20 nm em.; mCherry: 560 ± 20 nm ex.; 610 ± 20 nm em., optimal gain). For reporters with weak signals, 20 μl of bacterial cultures at the 6 h timepoint were diluted into 80 μl of M63 salts (KD Medical, Columbia, MD) in a SensoPlate Plus glass bottom microplate (Greiner Bio-One), and then cell fluorescence and optical density (OD_600_) were measured using the TECAN microplate reader with the same setting as above. By doing this, background fluorescence was minimized, and cell density was measured more precisely. To calculate fluorescence fold changes, fluorescence and OD_600_ readings from a well with medium blank were subtracted, and then fluorescence signals were normalized to cell density (OD_600_). Normalized fluorescence of each sample was further compared to that of the mean value of controls. Data were plotted as mean and standard deviation using GraphPad Prism 8. Unpaired two-tailed Student’s t-test was used to calculate statistical significance. Not significant (ns), P > 0.05; (*) P < 0.05; (**) P < 0.01; (***) P < 0.001; (****) P < 0.0001.

### Western blotting

Western blotting was carried out as previously described (63). Briefly, bacterial cells were pelleted, washed once with 1× PBS (phosphate-buffered saline), then directly resuspended in 1× SDS (sodium dodecyl sulphate) sample buffer (New England BioLabs) per 1 ml of OD_600_ of 1.0 cells in 200 μl sample buffer. 5-10 μl of boiled supernatant of whole cell lysates was resolved on 4-12% Bis-Tris NuPAGE protein gel (Invitrogen) in 1×MES buffer (Invitrogen) at 150 V for 1 h. Resolved protein samples were transferred onto a PVDF membrane using the iBlot 2 dry blotting system (ThermoFisherScientific). Incubation of the membrane with primary and secondary antibodies was done at room temperature for 1 h each. Before and after secondary antibodies, the blot was washed 3×10 min with PBS+0.1% Tween 20. Western signals were visualized using CDP-star substrate (Invitrogen) on a Bio-Rad ChemiDoc MP Imaging system. Antibodies were used at the following dilutions: mouse monoclonal anti-RFP antibody (MA5-15257) (1:3000; ThermoFisherScientific), monoclonal anti-FLAG M2-alkaline phosphatase antibody (1:3000; Sigma, A9469), monoclonal mouse anti-EF-Tu antibody (1:10,000; LSBio), polyclonal rabbit anti-Hfq antibodies (1:2500) (29), polyclonal rabbit anti-His antibodies (sc-803) (1:1000, Santa Cruz Biotechnology), anti-rabbit AP-conjugated IgG (1:5000; Cell Signaling Technology), and anti-mouse AP-conjugated IgG (1:10,000; Santa Cruz Biotechnology).

### Northern blotting

Total RNAs from bacterial cultures were prepared using the hot phenol method (52) with Phase Lock Gel (PLG Heavy, Quantabio), and RNA concentrations were determined using a Nanodrop 1000 spectrophotometer (ThermoScientific). To visualize RNAs of less than 500 nts, 3 μg of total RNAs of each sample were resolved on 10% Criterion TBE-urea polyacrylamide gel (Bio-Rad), and processed with the standard procedure described previously (63). To visualize mRNAs, 10 μg of total RNA were resolved on 1.2% agarose gels in 1×MOPS buffer, and processed with the procedure described previously (23). 5′-biotinylated Northern probes (see Table S3) were used, and the Brightstar BioDetect kit (Ambion) with the CDP-Star substrate was used for visualization of Northern signals using the Bio-Rad ChemiDoc MP Imaging system.

### Simultaneously mapping RNA 5′ and 3′ends by RNA self-ligation, reverse transcription (RT), PCR cloning and sequencing

To map the 5′ and 3′ ends of RyhB transcripts from cells expressing AspX or RNase T variants, a modified RACE (rapid amplification of cDNA ends) method (36) was used, which involves RNA self-ligation, reverse transcription, PCR cloning and sequencing of clones. Briefly, total RNAs from bacterial cultures at OD_600_ ∼2.0 were extracted. 3 μg of total RNA was treated with RQ1 DNase (Promega), and then purified by phenol chloroform extraction and ethanol precipitation. Purified RNAs were then treated with TAP (Tobacco Acid Pyrophosphatase, Epicentre) to convert 5′ ends of RNA to monophosphate, and the resultant RNAs were purified by phenol chloroform extraction and ethanol precipitation. After purification, RNAs were ligated by T4 RNA ligase 1 (New England BioLabs), and purified by ethanol precipitation. Purified ligated RNA samples were split evenly (one portion used for -RT control) for reverse transcription with *ryhB*-specific oligo JC413 using SuperScript III reverse transcriptase following the standard protocol. After reverse transcription, RNase H was added to remove the RNA templates, and 2 μl of cDNA products were used for PCR amplification with oligos JC413/JC414. Resultant PCR products were cloned into a medium copy number plasmid pHDB3 (15-20 copy/cell) through restriction cloning (EcoRI/ BamHI). 12 individual clones from each control group and 36 clones from treatment group were sequenced using the pBRlib_For primer. Sequenced hybrids were mapped to the *ryhB* gene, allowing determination of the 5′ and 3′ ends through junction sequences.

### Bacterial two hybrid (BACTH) assay

The BACTH system used in this study is based on reconstitution of adenylate cyclase CyaA activity in *E. coli* strain BTH101, which lacks a functional *cyaA* gene (64). The proteins YicC and PNPase were fused to the T18- and T25-fragments of the CyaA protein from *Bordetella pertussis*. The T18 and T25 domains interact to form a functional CyaA protein that synthesizes cAMP only when the fusion proteins interact with each other. cAMP-CRP drives *lacZ* reporter gene expression, and β -galactosidase activities indicate the strength of interaction between the fusion proteins. Fusions of YicC and PNPase to the T25 CyaA fragment were expressed from plasmids pKT25-*yicC* and pKT25-*pnp*, respectively; fusions of YicC and PNPase to the T18 CyaA fragment were expressed from plasmids pUT18-*yicC* and pUT18-*pnp*, respectively. Strain BTH101 was co-transformed with combinations of pUT18 and pKT25 plasmids expressing the T18- and T25-fusion proteins. To measure their interactions, the co-transformants were spotted onto LB plates containing X-Gal (5-Brom-4-chlor-3-indoxyl-β-D-galactopyranoside) (40 μg/ml), Kan and Amp. The plates were incubated at 30°C for ∼60 h. Colony blueness indicates β-galactosidase activity and thus interactions of the fusion proteins. To quantify the protein interactions, co-transformants were grown at 30°C for 16 h in LB containing Amp and Kan, and β-galactosidase activities were determined using the standard Miller assay.

## Acknowledgements

We thank Teppei Morita, Sandra Wolin, Gisela Storz, and members of the Gottesman lab for their comments on the manuscript, and Jay Zhu for the BACTH work conducted by Jiandong Chen in his lab in the University of Pennsylvania. This work was supported by the Intramural Research Program of the NIH, National Cancer Institute, Center for Cancer Research. Jiandong Chen acknowledges support from NCI Director’s Intramural Innovation Awards 479434 and 602936.

