## Supplemental figs, tables and methods for "A fluorescence-based genetic screen reveals diverse mechanisms silencing small RNA signaling in *E. coli*"

**This supplement contains:**

Supplementary Figures (S1-S12) with Legends
Supplementary Materials and Methods
Tables S1 to S3
Supplemental References

**TABLE OF CONTENTS**


**Figure S1** **A chromosomal fluorescent reporter system monitors Hfq/sRNA signaling through a tandem target mRNA fluorescence fusion in *E. coli***

**Figure S2** **Flowchart of fluorescence-based genomic suppressor screen for silencers of sRNA signaling**

**Figure S3** **Genomic library screen identified candidate regulators of reporter fusions**

**Figure S4** **Effects of individual factors on expression of additional mRNA targets of RyhB**

**Figure S5** **Chromosomal deletion of individual regulators showed modest effects on *sodB* reporter expression**

**Figure S6** **Effects from overproduction of individual regulators on control and *mutS* fusion expression**

**Figure S7** **Hfq-dependent sRNA CyaR, ChiX and McaS can compete for Hfq**

**Figure S8** **RNase T promotes RyhB sRNA decay**

**Figure S9** **Effects from overproduction of individual factors on Hfq accumulation**

**Figure S10** **AspX is a novel transcript expressed at the 3'end of *aspA* mRNA**

**Figure S11** **Effects of AspX variants on RyhB regulation**

**Figure S12** **YicC, a conserved protein, selectively degrades RyhB sRNA**

**Supplementary Materials and Methods**

**Table S1** Strains used in this study

**Table S2** Plasmids used in this study

**Table S3** Primers, probes and synthetic gene fragments used in this study

### Figure S1

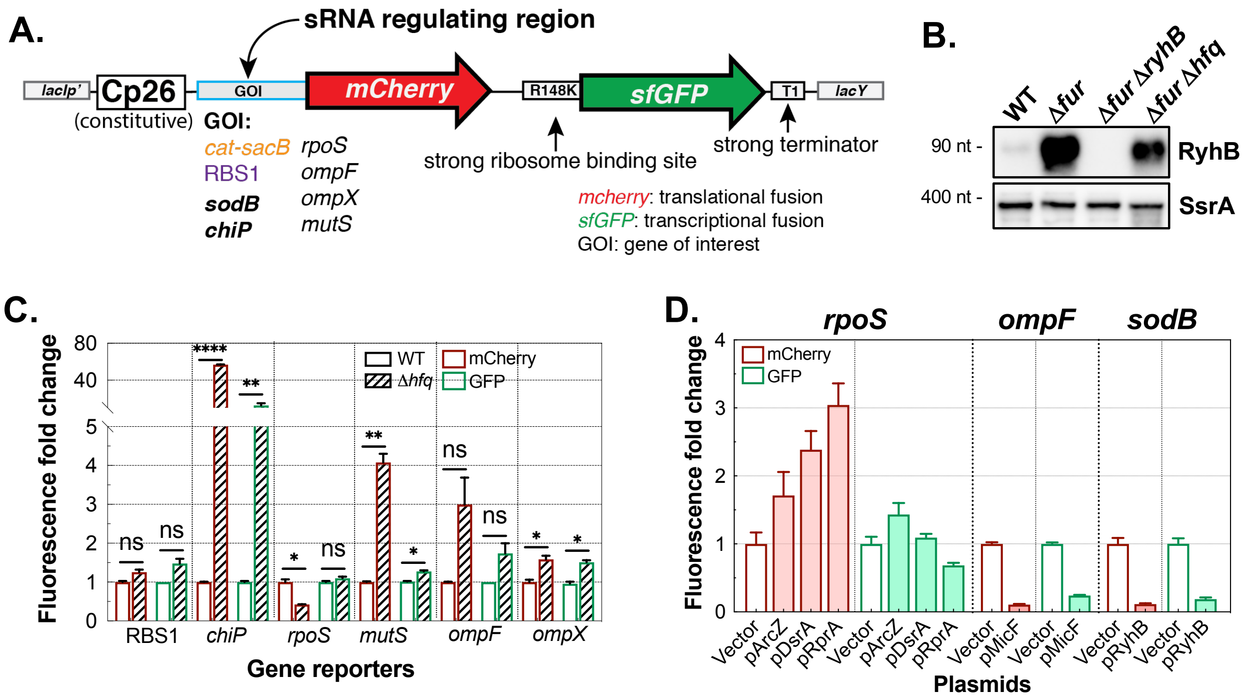


**Figure S1. A chromosomal fluorescent reporter system monitors Hfq/sRNA signaling through a tandem target mRNA fluorescence fusion in *E. coli***

**A**. Schematic of a tandem fluorescence reporter system designed for monitoring sRNA regulation of genes in *E. coli*. An operon consisting of a constitutive promoter (Cp26), the leader and initial translated region of a gene of interest (GOI) translationally fused to the mCherry gene and transcriptionally fused to a downstream superfolder GFP, ending with a ribosomal transcription terminator (T1), was engineered into the *E. coli* chromosome at the *lacI-lacZ* locus. GFP is translated from a potent ribosome binding sequence R148K (ACACAGAAAGCAAAAACATTAAGGGGGTAATA) (1). Fluorescent gene translational fusions were made by $\lambda$Red recombineering of the leader and initial translated region of the GOI in a parental strain that harbors a *cat-sacB* cassette upstream of the mCherry gene. All constructions were verified by PCR and sequencing.

**B**. RyhB levels modulated by Fur and Hfq. WT (JC1246), Δ*fur* (JC1248), Δ*fur*Δ*ryhB* (JC1252), Δ*fur*Δ*hfq* (JC1269) were grown to $OD_{600}$ of ~1.0 in LB, and RyhB levels were measured by Northern blot analysis. SsrA served as a loading control.

**C**. Tests of Hfq regulation of multiple fusions. WT or Δ*hfq* strains of mRNA reporter *chiP* (JC1247 [WT], JC1267 [Δ*hfq*]), *rpoS* (JC1197 [WT], JC1264 [Δ*hfq*]), *mutS* (JC1286 [WT], JC1297 [Δ*hfq*]), *ompF* (JC1198 [WT], JC1265 [Δ*hfq*]), *ompX* (JC1285 [WT], JC1296 [Δ*hfq*]) or the synthetic RBS1 control (GCAATTTTAAGGAGGTAACT) (JC1245 [WT], JC1263 [Δ*hfq*]) were grown in LB in a 96-well microplate for 6 h at 37°C before measuring cell optical density ($OD_{600}$) and mCherry/GFP fluorescence using a TECAN Spark 10M microplate reader. Biological duplicates of each sample were measured. Unpaired two-tailed Student's t-test was used to calculate statistical significance. Not significant (ns), $P > 0.05$; (∗) $P < 0.05$; (∗∗) $P < 0.01$; (∗∗∗∗) $P < 0.0001$.

**D**. Multicopy sRNA regulation of mRNA fusions. Reporter strains of *rpoS* (JC1329), *ompF* (JC1330) or *sodB* (JC1246) transformed with plasmids expressing ArcZ, DsrA, RprA; MicF or RyhB sRNA as listed, were grown in LB+Amp+IPTG (50 μM) for 5 h at 37°C before measuring cell optical density ($OD_{600}$) and fluorescence. Relative fluorescence was normalized to the empty vector control of each fusion. 3-6 replicates were assayed for each strain, and data are presented as mean with standard deviation.

**Figure S2**

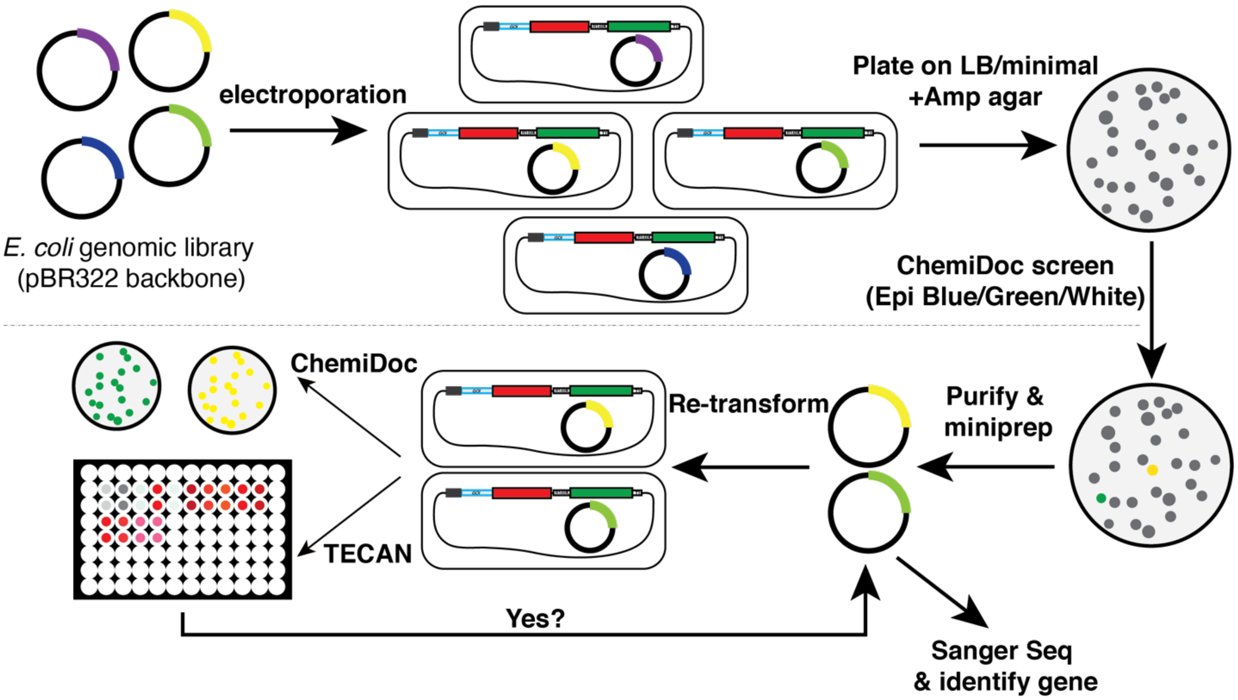


**Figure S2. Flowchart of fluorescence-based genomic suppressor screen for silencers of sRNA signaling**

A plasmid library consisting of 1.5 – 5 kb *E.coli* genomic fragments (2) in a plasmid encoding ampicillin resistance was transformed into a fluorescent reporter strain, in which changes in fluorescence provides a readout of sRNA activity. For the screens carried out here, sRNA regulation led to low fluorescence; colonies were screened for fragments that interfered with sRNA function and thus had increased fluorescence. Transformants were plated on selective LB ampicillin or M63 glucose ampicillin agar plates (50 plates, ~500 CFUs/plate), and screened for clones expressing elevated fluorescence, using the Bio-Rad ChemiDoc MP Imaging system. Candidate clones were purified 2-4 times, and then plasmids were extracted and re-transformed into the parental reporter strain for verification, both qualitatively on agar plates and quantitively in liquid culture. Plasmids from verified positive clones were sequenced from the flanking sequence of the insert using sequencing oligos pBRlib_For and pBRlib_Rev, and sequencing data were mapped to the *E. coli* K-12 genome using EcoCyc BLASTn search. All purified positive clones remained positive after retransformation. Bacteria grown on minimal agar had lower background fluorescence, and thus minimal agar plates may be preferred for identifying clones showing modest increases of fluorescence compared to the parental control.

### Figure S3

**A.**

| Summary of genomic library screen results | | | |
|---|---|---|---|
| *Reporter* | *# clones screened* | *# positive clones* | *Positive rate* |
| ***chiP*** | 45,000 | 25 | 0.0006 |
| ***sodB*** | 17,800 | 26 | 0.0015 |

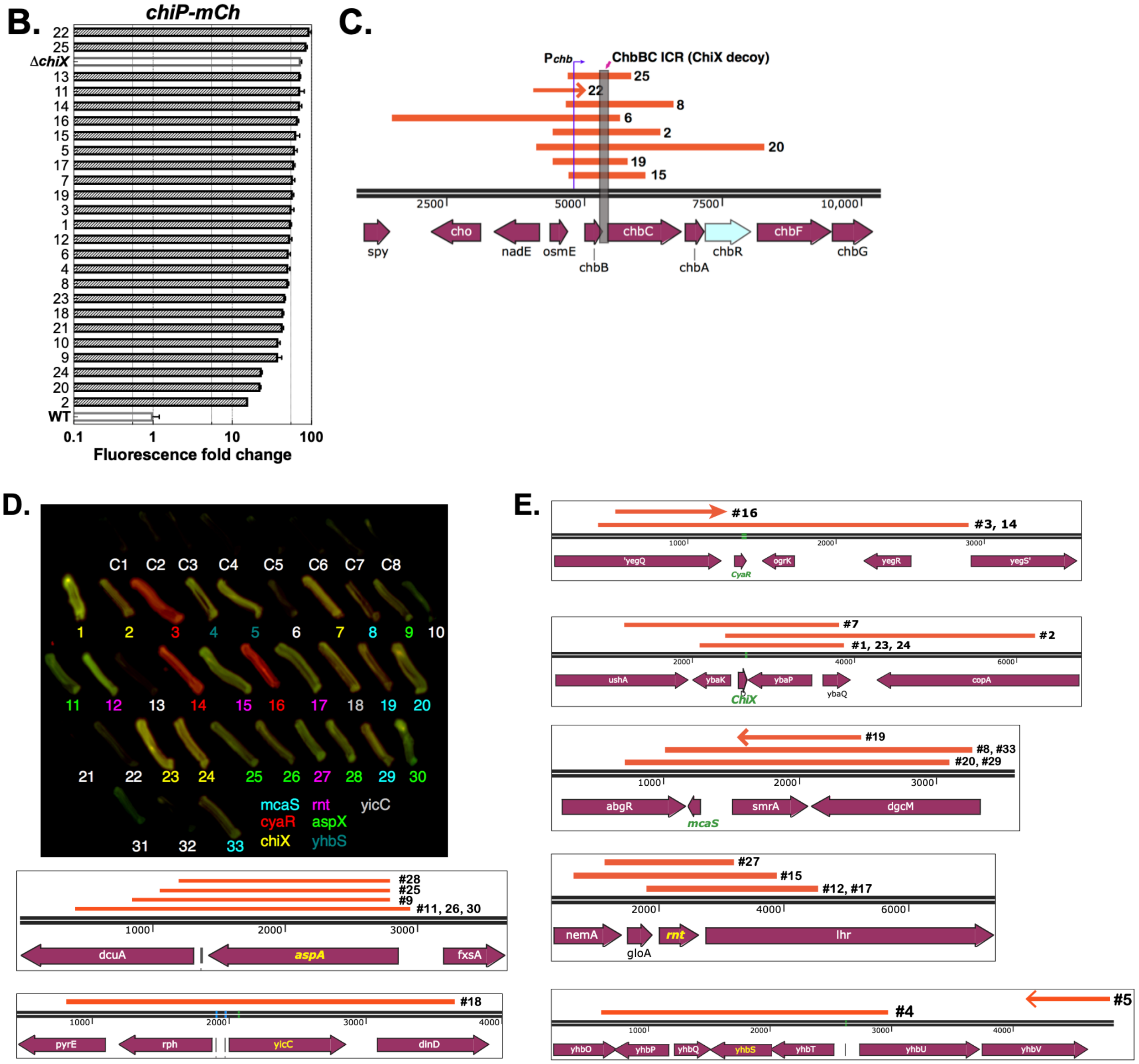

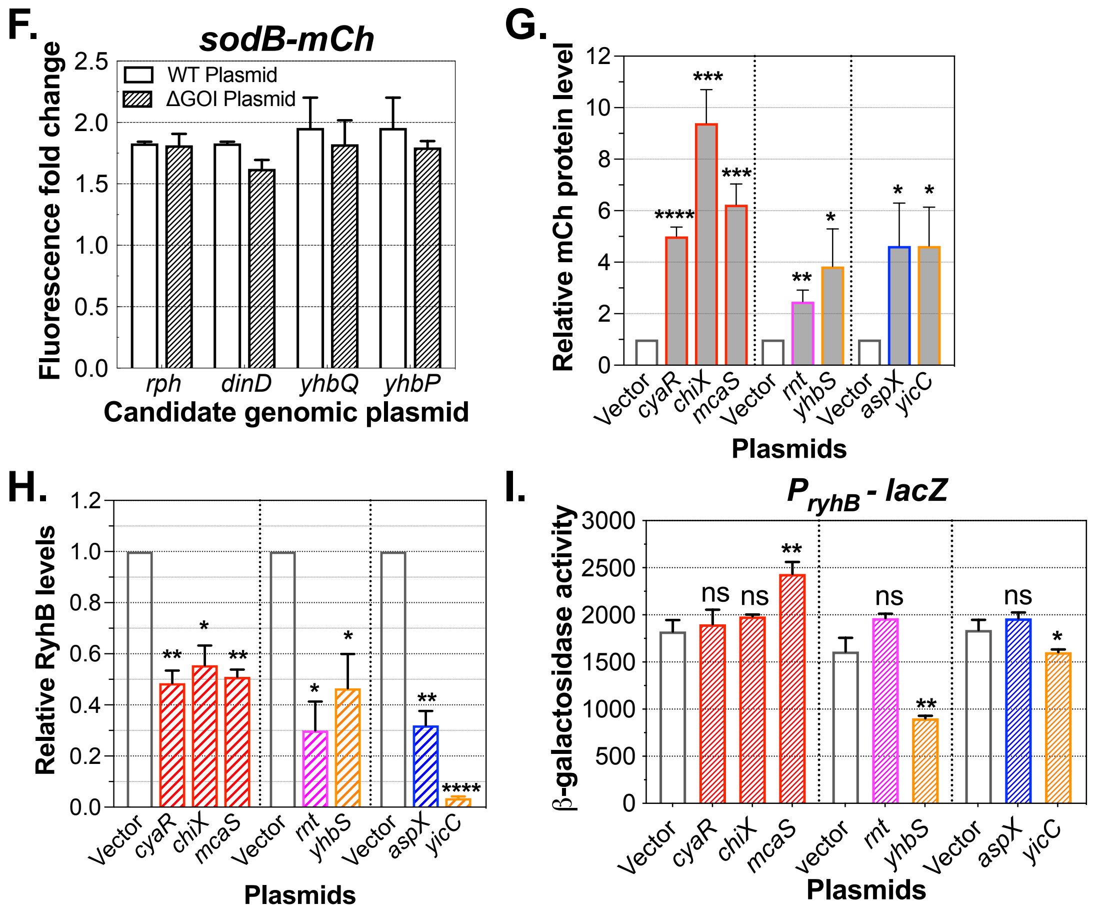


**Figure S3. Genomic library screen identified candidate regulators of reporter fusions**

**A**. Summary of genomic library screen for regulators of sRNA signaling: ChiX (*chiP* fusion) and RyhB (*sodB* fusion). 45,000 and 17,800 total clones were screened on the *chiP* and *sodB* fusions, representing ~29× and ~12× coverage of the *E. coli* genome, respectively.

**B**. Fluorescence of WT (JC1200, parental strain expressing basal fluorescence), Δ*chiX* (JC1244, positive control) and 25 high fluorescence clones identified from genomic library screen of the *chiP* fusion. Data are plotted on a log10 scale.

**C**. Plasmids from eight positive clones expressing different levels of ChiP-mCherry fluorescence were sequenced and mapped to the *E. coli* K-12 genome. All eight inserts (orange bars) mapped to the same *chbBCARFG* locus, where a ChiX decoy, the *chbBC* intercistronic region (ICR), is encoded (gray box) (3). Occasionally, the sequence of the two ends of the insert were mapped to two distant regions of the chromosome, presumably representing two inserts in one vector. In this case, only the relevant region is shown as an arrow. The arrow covers the sequenced region (~1 kb) and the arrowhead shows the direction of the sequence. The vertical line and L-shaped arrow at the top of the orange bars represents the promoter of the *chbBCARFG* operon.

**D**. Control (C1-C8) and candidate clones (#1-33) from the *sodB* fusion screen were patched on LB+Amp agar and imaged for mCherry and GFP using the Bio-Rad ChemiDoc MP Imaging system. C1-C8 are clones selected from the original screen plates that expressed the low basal fluorescence (not visible in figure). 7 out of 33 clones selected for higher fluorescence (white

numbers) showed insignificant increases of fluorescence and were left out of further analysis. Numbers of other clones are color-coded to reflect the region they encoded (see gene names at bottom of panel).
**E**. Plasmids of 26 positive clones were sequenced and mapped to the *E. coli* K-12 genome using BLASTn search (https://ecocyc.org/ECOLI/blast.html). The inserts (orange bars) clustered to seven distinct genomic loci and are numbered as in **Fig. S3D**. Occasionally, two ends of the insert were mapped to two distant regions, presumably representing two inserts into one vector; in this case, only the relevant region is shown as an arrow.
**F**. Identification of relevant gene in two genomic regions. Plasmids containing deletions of genes (ΔGOI Plasmid), as indicated, within plasmid #18 (*yicC* clone) or #4 (*yhbS* clone) and the parental plasmids were introduced into the *sodB* reporter strain (JC1248) and grown for 6 h at 37°C in LB+Amp before measuring $OD_{600}$ and fluorescence. Fluorescence fold change is relative to the mean of controls (C1-C3).
**G**. Quantitation of mCherry Western blot signals in **Fig. 3D** using Image Studio Lite (LI-COR Biosciences) from two independent experiments.
**H.** Quantitation of RyhB Northern blot signals in **Fig. 3E** using Image Studio Lite (LI-COR Biosciences) from two independent experiments. RyhB in pAspX samples appear as a doublet, and both bands were included in the bar shown. RyhB signals were normalized to the SsrA loading control.
**I**. Test of plasmid effects on the *ryhB* promoter. A strain carrying the *ryhB* promoter transcriptionally fused to *lacZ* fusion (JC1327; Δ*fur* to mimic situation in original fusion screening strain) was transformed with the pBRplac plasmid expressing CyaR, ChiX or McaS, or the pQE-80L plasmid expressing RNase T or YhbS, or the pBR* plasmid expressing AspX or YicC. Transformed cells were grown for 5 h in LB supplemented with appropriate antibiotics and IPTG (50μM) and β-galactosidase activities were measured from biological triplicates.
Data are presented as mean with standard deviation. Unpaired two-tailed Student's t-test was used to calculate statistical significance. Not significant (ns): $P > 0.05$; (∗): $P < 0.05$; (∗∗): $P < 0.01$; (∗∗∗): $P < 0.001$; (∗∗∗∗): $P < 0.0001$.

**Figure S4**

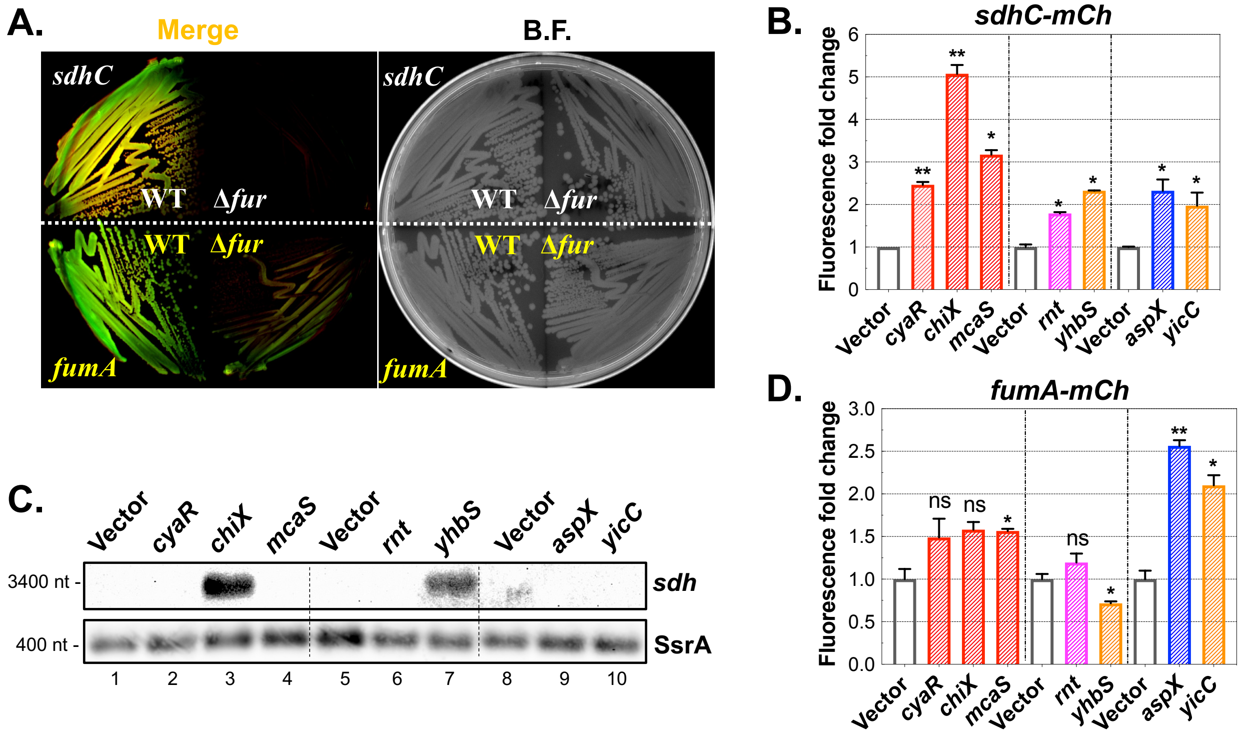


**Figure S4. Effects of individual factors on expression of additional mRNA targets of RyhB**

**A**. Tandem fluorescent reporter strain for *sdhC* (WT: JC1334; Δ*fur*: JC1339) or *fumA* (WT: JC1335; Δ*fur*: JC1340) were streaked on LB agar and imaged after 16 h growth at 37°C using the Bio-Rad ChemiDoc MP Imaging system. Right image shows bright field (B.F.) image.

**B**. Reporter strains for *sdhC* (JC1339) overexpressing CyaR, ChiX or McaS sRNA from pBR*plac*, or His-RNase T or His-YhbS from pQE-80L, or AspX or YicC from pBR* were grown in 96-well microplates with LB IPTG (50 μM) and appropriate antibiotics for 6 h at 37°C. Fluorescence for biological duplicates of each sample were measured at the endpoint using the TECAN Spark 10M microplate reader.

**C**. Endogenous *sdhCDAB* mRNAs were measured by Northern blotting using total RNA samples used in **Fig. 3E**.

**D**. Reporter strains for *fumA* (JC1340) overexpressing CyaR, ChiX or McaS sRNA from pBR*plac*, or RNase T, YhbS from pQE-80L, or AspX, YicC from pBR* were grown in 96-well microplates with LB IPTG (50 μM) and appropriate antibiotics for 6 h at 37°C. Fluorescence for biological duplicates of each sample were measured at the endpoint using the TECAN Spark 10M microplate reader.

Data are presented as mean with standard deviation. Unpaired two-tailed Student's t-test was used to calculate statistical significance. Not significant (ns): $P > 0.05$; (∗): $P < 0.05$; (∗∗): $P < 0.01$.

**Figure S5**

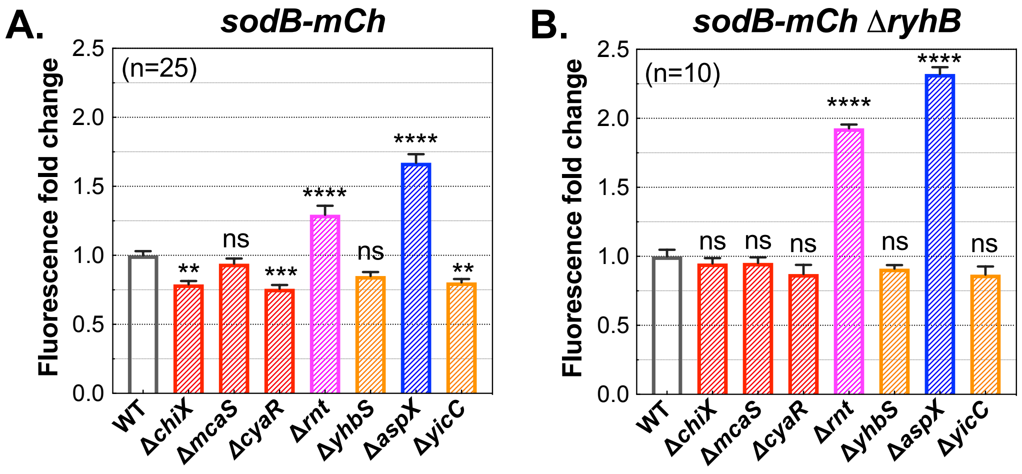


**Figure S5. Chromosomal deletion of individual regulators showed modest effects on *sodB* reporter expression**

**A**. WT (JC1246) or mutants deleted of chromosomal *chiX* (JC1355), *mcaS* (JC1356), *cyaR* (JC1357), *rnt* (JC1358), *yhbS* (JC1359), *aspX* (JC1360) or *yicC* (JC1361) were grown in LB in a microplate for 6 h at 37°C prior to fluorescence measurement.

**B**. A *ryhB* null mutation was introduced to the same set of strains (JC1270, JC1367-JC1373) as in **Fig. S5A** and fluorescence was measured after growth in the same conditions as above.

Δ*rnt* and Δ*aspX* strains exhibited slow growth in LB. 10 or more biological repeats were measured for each strain, and data are plotted as mean and standard deviation.

Unpaired two-tailed Student's t-test was used to calculate statistical significance. Not significant (ns): P > 0.05; (∗∗): P < 0.01; (∗∗∗): P < 0.001; (∗∗∗∗): P < 0.0001.

**Figure S6**

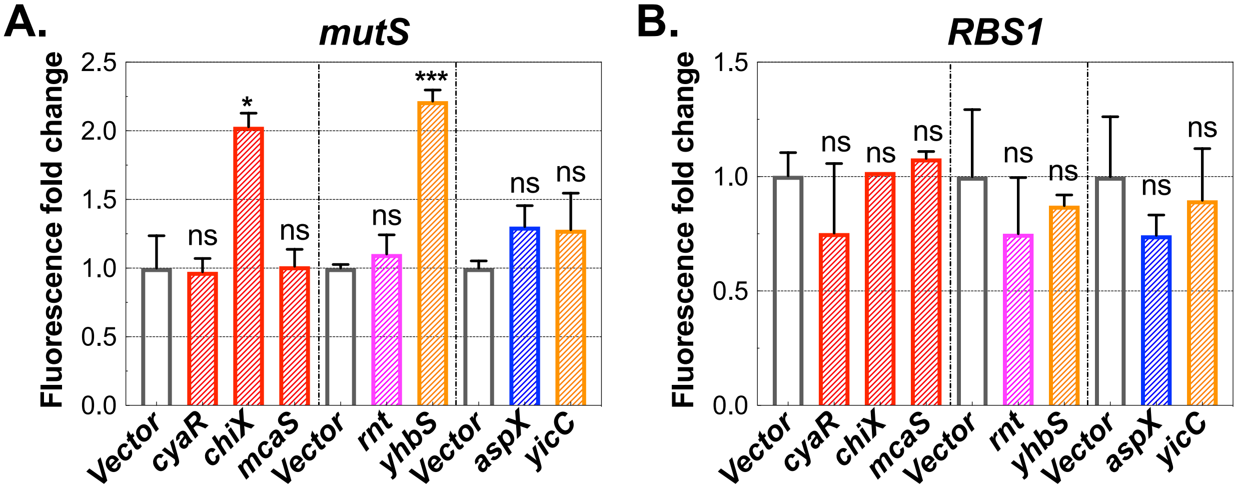


**C.**

Relative quantitation of CyaR and ArcZ sRNA from Northern blots in Fig. 4C

| | *Vector* | *cyaR* | *chiX* | *mcaS* | *Vector* | *rnt* | *yhbS* | *vector* | *aspX* | *yicC* |
|---|---|---|---|---|---|---|---|---|---|---|
| **CyaR** | 1.00 | 10.72 | 0.22 | 0.07 | 1.00 | 0.20 | 0.05 | 1.00 | 1.91 | 1.88 |
| **ArcZ** | 1.00 | 0.65 | 0.79 | 0.55 | 1.00 | 0.57 | 0.46 | 1.00 | 1.70 | 1.74 |

**Figure S6. Effects from overproduction of individual regulators on control and *mutS* fusion expression**

**A**, **B**. Reporter strains for *mutS* (JC1332) or RBS1 (JC1245) overexpressing CyaR, ChiX or McaS sRNA from pBRplac, or His-RNase T, His-YhbS from pQE-80L, or AspX, YicC from pBR* were measured for mCherry fluorescence after 6 h induction at 37°C using a TECAN microplate reader. A final concentration of 50μM of IPTG was added at the time of inoculation into the microplates. Biological triplicates of each sample were measured. Unpaired two-tailed Student's t-test was used to calculate statistical significance. Not significant (ns): P > 0.05; (∗): P < 0.05; (∗∗∗): P < 0.001.

**C**. Quantitation of Northern blot signals for CyaR and ArcZ in **Fig. 4C**. Relative sRNA levels were calculated by normalization to the SsrA loading control first and then further compared to that of the vector control in each group.

**Figure S7**

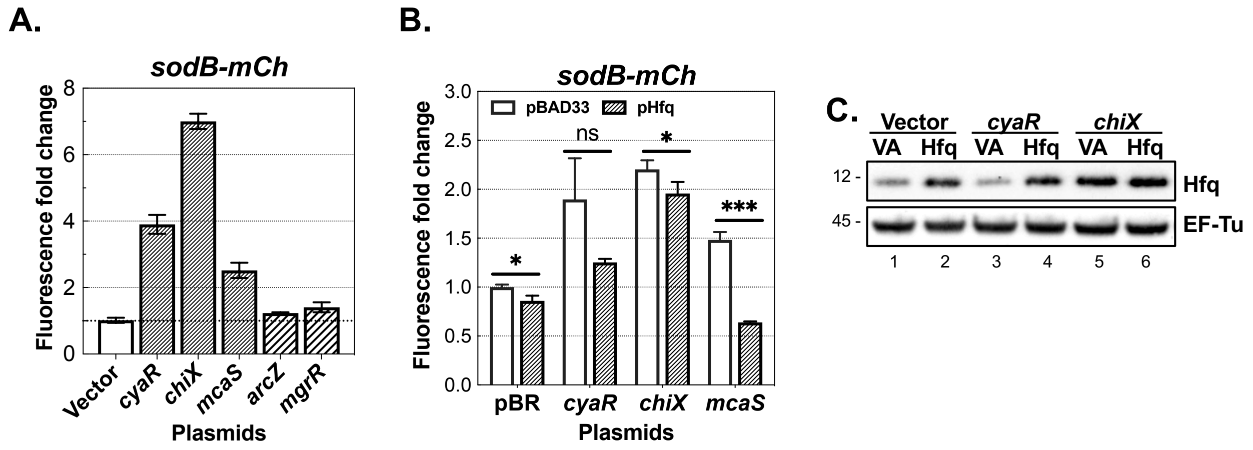


**Figure S7. Hfq-dependent sRNA CyaR, ChiX and McaS can compete for Hfq**

**A**. The *sodB* reporter strain (JC1249) expressing sRNA CyaR, ChiX, McaS, ArcZ or MgrR from the pBRplac plasmid were grown for 6 h at 37°C in microplates with LB+Amp+IPTG (50 μM) prior to measuring mCherry fluorescence.

**B**. *sodB* reporter strains (JC1322) expressing sRNA CyaR, ChiX, or McaS from the pBRplac plasmid and Hfq from pBAD33 were grown for 15 h at 37°C in LB+Amp+Chl+IPTG (50 μM)+ arabinose (0.1%) before measuring mCherry fluorescence. Relative fluorescence of each strain was calculated by normalizing to the control harboring two empty vectors. Biological triplicates of each sample were measured. Unpaired two-tailed Student's t-test was used to calculate statistical significance. Not significant (ns): $P > 0.05$; (*): $P < 0.05$; (***): $P < 0.001$.

**C**. Whole cell lysates prepared from the cells in **Fig. S7B** were analyzed by western blot for Hfq. EF-Tu served as a loading control. Note, due to poor recovery of samples coexpressing McaS and Hfq, these samples were not included for this analysis.

**Figure S8**

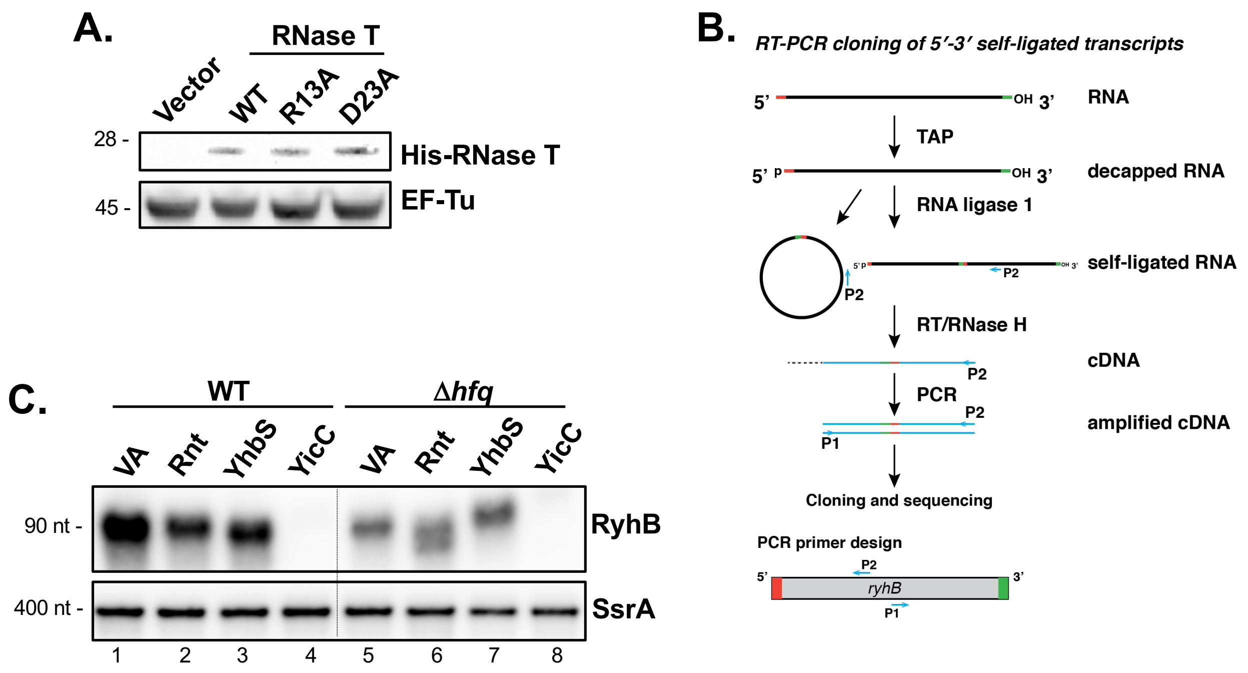


**Figure S8. RNase T promotes RyhB sRNA decay**

**A**. Whole cell lysates were prepared from cultures in **Fig. 5A** and analyzed by Western blot using anti-His and anti-EF-Tu antibodies.

**B**. Schematic diagram for simultaneously mapping RNA 5' and 3' ends using RNA self-ligation, reverse transcription, PCR cloning and sequencing (4). Outward-facing oligos P1 and P2 are shown, which do not make an amplicon for non-ligated products.

**C**. WT (JC1341) or Δ*hfq* strain (JC1352) harboring vector (pQE-80L) or plasmids expressing RNase T, YhbS or YicC plasmid were grown in LB+Amp for 2 h ($OD_{600}$ ~0.2) at 37°C, then induced with 100 μM IPTG for 2 additional hours. RyhB was measured by Northern blot analysis, and SsrA served as a loading control.

**Figure S9**

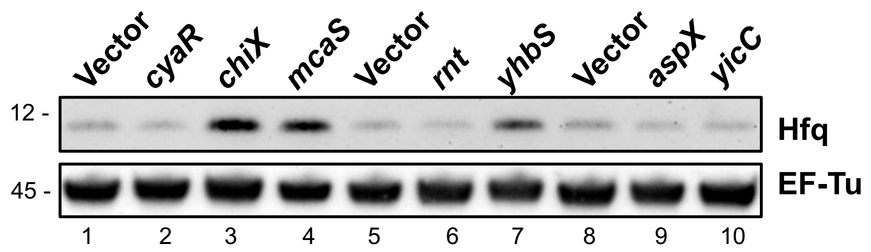


**Figure S9. Effects from overproduction of individual factors on Hfq accumulation**
*sodB* reporter strain (JC1318) overexpressing CyaR, ChiX or McaS sRNA from the pBRplac plasmid, or His-tagged RNase T, YhbS from the pQE-80L plasmid, or AspX, YicC from the pBR* plasmid were induced with 50 μM IPTG for 6 h at 37°C; whole cell lysates were prepared for Western blot analysis of Hfq.

### Figure S10

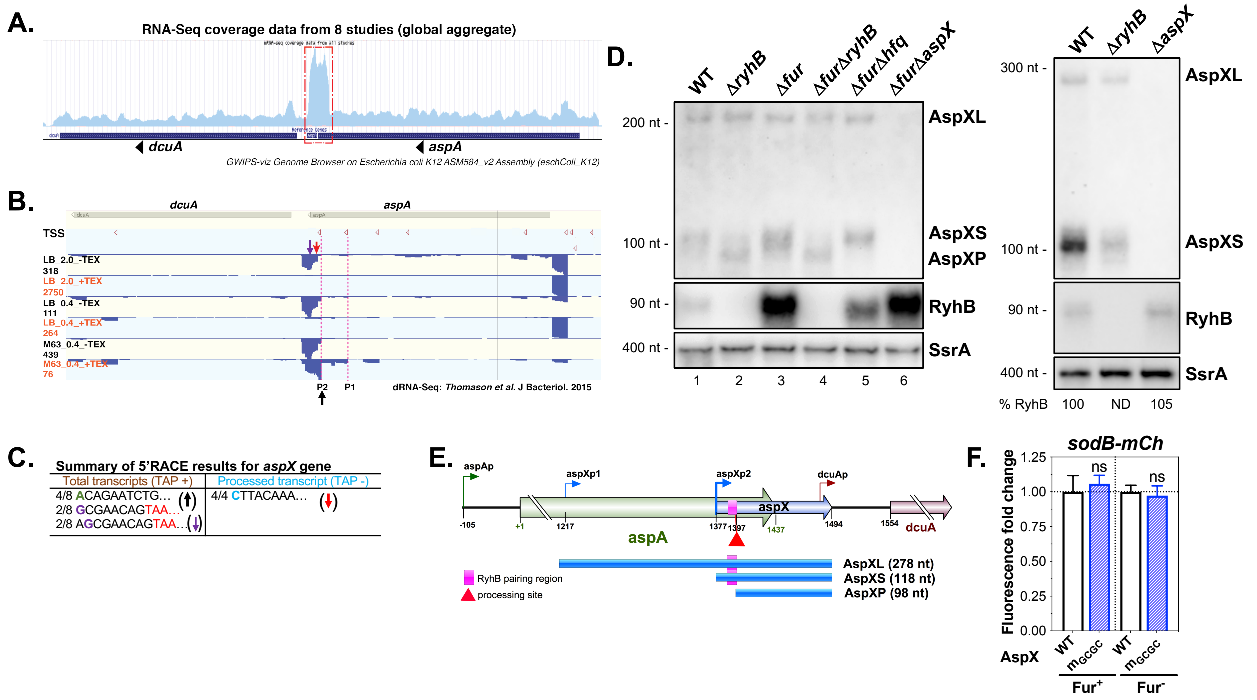

**Figure S10. AspX is a novel transcript expressed at the 3'end of *aspA* mRNA**

**A**. RNA-seq reads density of the *aspA-dcuA* region displayed in the GWIPS-viz Genome Browser (https://gwips.ucc.ie/index.html). The red dashed box highlights a putative new transcript at the 3'end of the *aspA* gene.

**B**. dRNA-Seq reads density of the *E. coli* K-12 MG1655 *aspA-dcuA* region (5). Two predominant transcription start sites (P1 and P2) and a cleavage site (red down arrow) for *aspX* are highlighted.

**C**. 5' RACE to map AspX 5' ends. *E. coli* MG1655 were grown in MOPS glucose medium to $OD_{600}$ of 1.0, and total RNAs were prepared for 5'RACE analysis (-/+TAP). Mapped transcription start sites (TSSs) are shown as the first nucleotide of each sequence. In the +TAP samples (both primary and processed transcripts can be detected), 4/8 sequences started with the same adenine (A), corresponding to the transcription start site of AspXS in Fig. S10B (P2, black upward arrow), and the other sequences started with either G or A nucleotide that precedes the G, corresponding to +51 and +50 respectively relative to the P2 start (denoted as a purple upward arrow in Fig. S10B). The TAA in red is the stop codon of the *aspA* gene. In the -TAP samples where only processed transcripts can be detected, all sequences started with the same cytosine (C), corresponding to the cleavage site in Fig. S10B (red down arrow, +21 relative to AspXS start).

**D**. WT (JC1246), Δ*ryhB* (JC1270), Δ*fur* (JC1248), Δ*fur*Δ*ryhB* (JC1252), Δ*fur*Δ*hfq* (JC1269), Δ*fur*Δ*aspX* (JC1319) and Δ*aspX* (JC1360) were grown in LB to $OD_{600}$ of 1.0, and RyhB and AspX were measured by Northern blot analysis. SsrA served as a loading control.

**E**. Schematic of the genomic context of *aspX*. The blue arrows indicate transcription start sites (TSS) for AspX, the purple box indicates the region that base pairs with RyhB, and the red arrowhead indicates a processing site for generating AspXP. Annotated TSSs of *aspA* and *dcuA* are shown as green and red L-shaped arrows, respectively.
**F**. *aspX*_WT (JC1374) or *aspX_m*$_{GCGC}$ chromosomal mutant (JC1375) in *fur*$^{+}$ strains or *aspX*_WT (JC1376) or *aspX_m*$_{GCGC}$ chromosomal mutant (JC1377) in *fur*$^{-}$ strains were grown in LB in a microplate for 6 h at 37°C prior to $OD_{600}$ and fluorescence measurements.
Twelve biological repeats were measured for each strain, and data are plotted relative to the value for the WT strain, as mean and standard deviation using GraphPad Prism 9.1.0.
Unpaired two-tailed Student's t-test was used to calculate statistical significance. Not significant (ns).

**Figure S11**

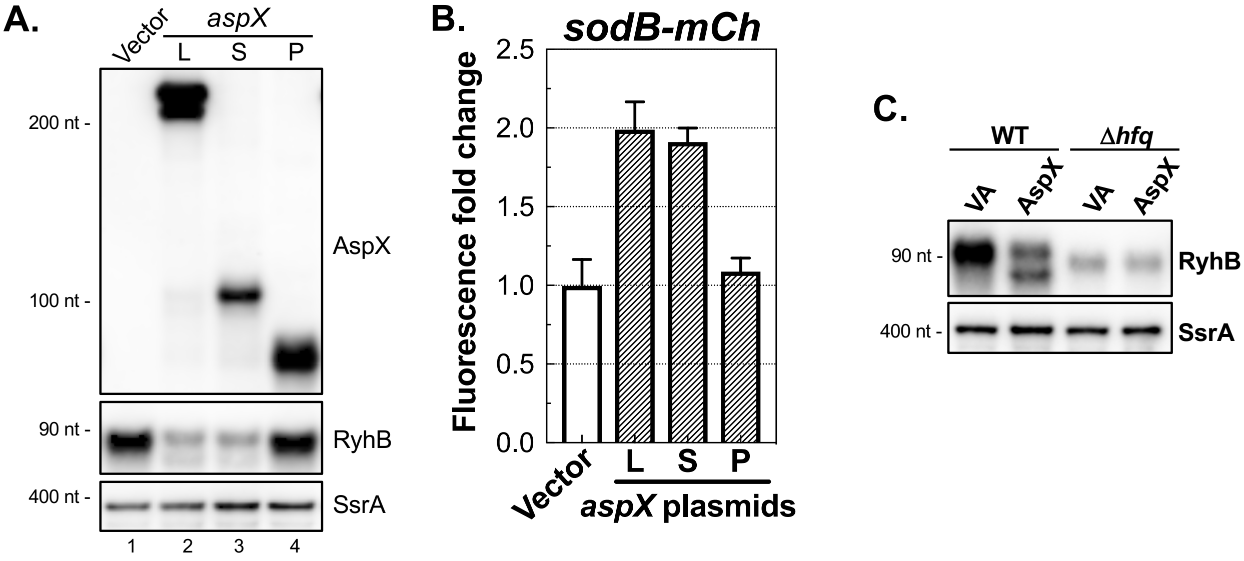


**Figure S11. Effects of AspX variants on RyhB regulation.**

**A**. The *sodB* reporter strain (JC1249) harboring AspX variant (L, S and P) plasmids were grown for 6 h at 37°C in LB+Kan+IPTG (50 μM). Total RNAs were extracted for Northern blot analysis of AspX, RyhB and SsrA.

**B**. The fluorescence normalized for $OD_{600}$ for cultures in **Fig. S11A** was determined using the TECAN microplate reader.

**C**. WT (JC1341) or Δ*hfq* (JC1352) harboring the vector or pAspX plasmid were grown in LB+Kan for 2 h ($OD_{600}$ ~0.2) at 37°C, then induced with 100 μM IPTG for 2 additional hours. RyhB and SsrA were probed for Northern blot analysis.

### Figure S12

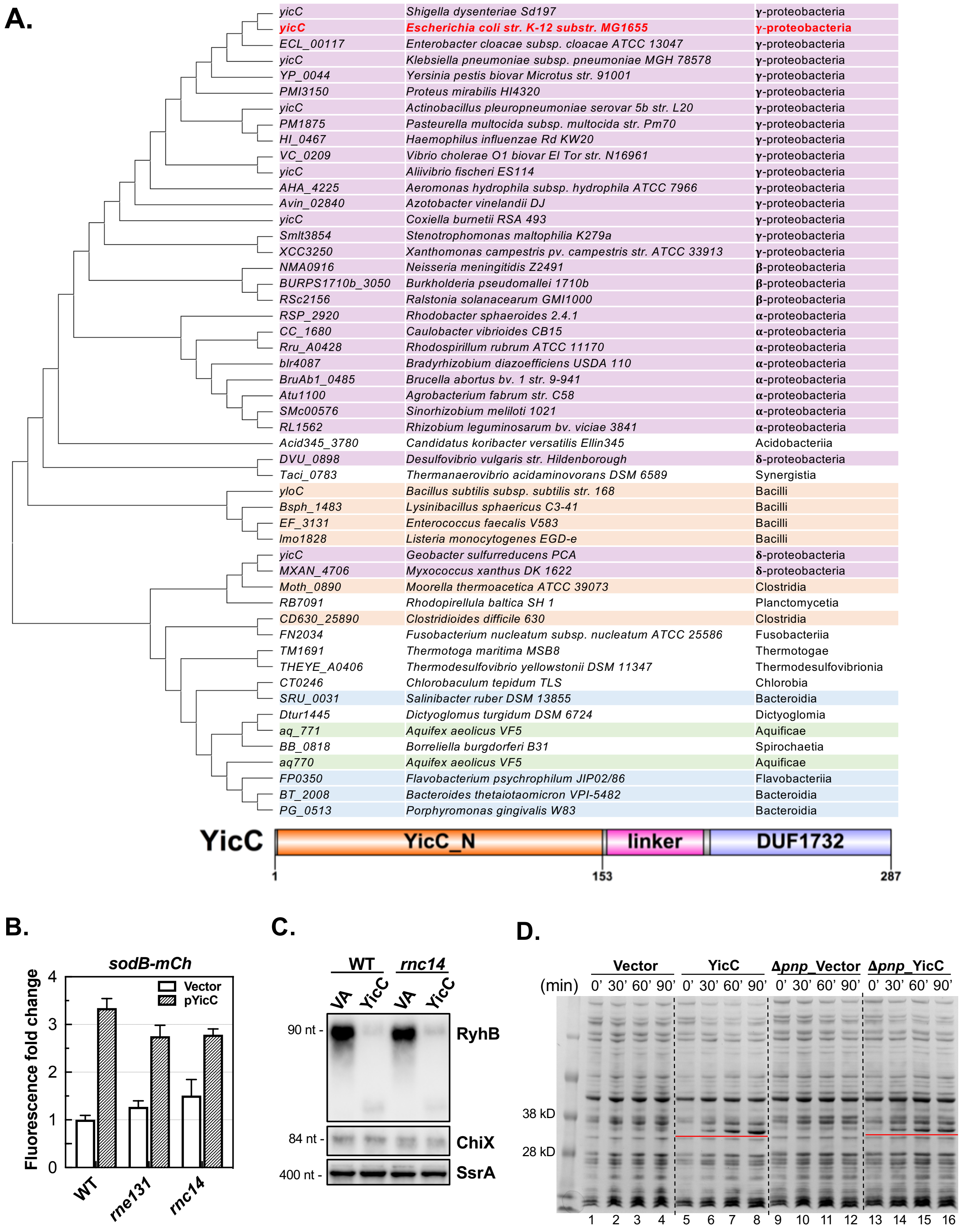

A.
yicC Shigella dysenteriae Sd197 γ-proteobacteria
yicC Escherichia coli str. K-12 substr. MG1655 γ-proteobacteria
ECL_00117 Enterobacter cloacae subsp. cloacae ATCC 13047 γ-proteobacteria
yicC Klebsiella pneumoniae subsp. pneumoniae MGH 78578 γ-proteobacteria
YP_0044 Yersinia pestis biovar Microtus str. 91001 γ-proteobacteria
PMI3150 Proteus mirabilis HI4320 γ-proteobacteria
yicC Actinobacillus pleuropneumoniae serovar 5b str. L20 γ-proteobacteria
PM1875 Pasteurella multocida subsp. multocida str. Pm70 γ-proteobacteria
HI_0467 Haemophilus influenzae Rd KW20 γ-proteobacteria
VC_0209 Vibrio cholerae O1 biovar El Tor str. N16961 γ-proteobacteria
yicC Aliivibrio fischeri ES114 γ-proteobacteria
AHA_4225 Aeromonas hydrophila subsp. hydrophila ATCC 7966 γ-proteobacteria
Avin_02840 Azotobacter vinelandii DJ γ-proteobacteria
yicC Coxiella burnetii RSA 493 γ-proteobacteria
Smlt3854 Stenotrophomonas maltophilia K279a γ-proteobacteria
XCC3250 Xanthomonas campestris pv. campestris str. ATCC 33913 γ-proteobacteria
NMA0916 Neisseria meningitidis Z2491 β-proteobacteria
BURPS1710b_3050 Burkholderia pseudomallei 1710b β-proteobacteria
RSc2156 Ralstonia solanacearum GMI1000 β-proteobacteria
RSP_2920 Rhodobacter sphaeroides 2.4.1 α-proteobacteria
CC_1680 Caulobacter vibrioides CB15 α-proteobacteria
Rru_A0428 Rhodospirillum rubrum ATCC 11170 α-proteobacteria
blr4087 Bradyrhizobium diazoefficiens USDA 110 α-proteobacteria
BruAb1_0485 Brucella abortus bv. 1 str. 9-941 α-proteobacteria
Atu1100 Agrobacterium fabrum str. C58 α-proteobacteria
SMc00576 Sinorhizobium meliloti 1021 α-proteobacteria
RL1562 Rhizobium leguminosarum bv. viciae 3841 α-proteobacteria
Acid345_3780 Candidatus koribacter versatilis Ellin345 Acidobacteriia
DVU_0898 Desulfovibrio vulgaris str. Hildenborough δ-proteobacteria
Taci_0783 Thermanaerovibrio acidaminovorans DSM 6589 Synergistia
yloC Bacillus subtilis subsp. subtilis str. 168 Bacilli
Bsph_1483 Lysinibacillus sphaericus C3-41 Bacilli
EF_3131 Enterococcus faecalis V583 Bacilli
lmo1828 Listeria monocytogenes EGD-e Bacilli
yicC Geobacter sulfurreducens PCA δ-proteobacteria
MXAN_4706 Myxococcus xanthus DK 1622 δ-proteobacteria
Moth_0890 Moorella thermoacetica ATCC 39073 Clostridia
RB7091 Rhodopirellula baltica SH 1 Planctomycetia
CD630_25890 Clostridioides difficile 630 Clostridia
FN2034 Fusobacterium nucleatum subsp. nucleatum ATCC 25586 Fusobacteriia
TM1691 Thermotoga maritima MSB8 Thermotogae
THEYE_A0406 Thermodesulfovibrio yellowstonii DSM 11347 Thermodesulfovibrionia
CT0246 Chlorobaculum tepidum TLS Chlorobia
SRU_0031 Salinibacter ruber DSM 13855 Bacteroidia
Dtur1445 Dictyoglomus turgidum DSM 6724 Dictyoglomia
aq_771 Aquifex aeolicus VF5 Aquificae
BB_0818 Borreliella burgdorferi B31 Spirochaetia
aq770 Aquifex aeolicus VF5 Aquificae
FP0350 Flavobacterium psychrophilum JIP02/86 Flavobacteriia
BT_2008 Bacteroides thetaiotaomicron VPI-5482 Bacteroidia
PG_0513 Porphyromonas gingivalis W83 Bacteroidia
YicC
YicC_N
linker
DUF1732
1
153
287
B.
sodB-mCh
Fluorescence fold change
0
1
2
3
4
Vector
pYicC
WT
rne131
rnc14
C.
WT
rnc14
VA
YicC
VA
YicC
90 nt -
RyhB
84 nt -
ChiX
400 nt -
SsrA
D.
Vector
YicC
Δpnp_Vector
Δpnp_YicC
(min)
0' 30' 60' 90' 0' 30' 60' 90' 0' 30' 60' 90' 0' 30' 60' 90'
38 kD
28 kD
1 2 3 4 5 6 7 8 9 10 11 12 13 14 15 16

E.

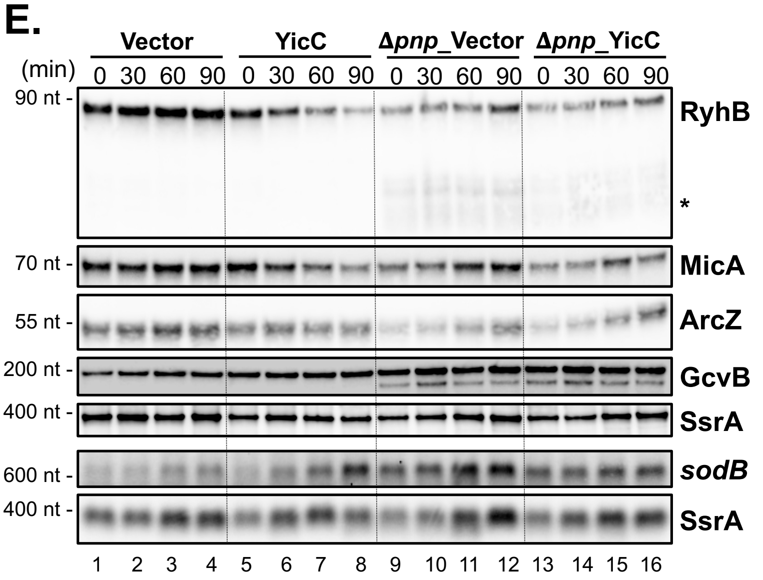


F.

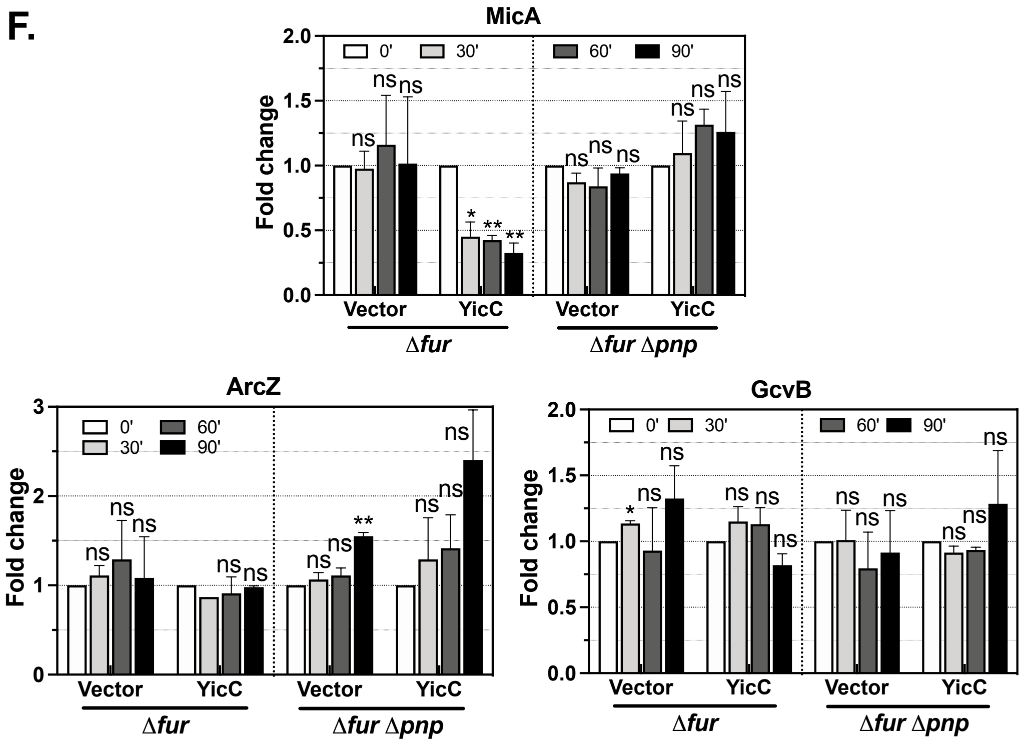


**Figure S12. YicC, a conserved protein, selectively degrades RyhB sRNA.**

**A**. Evolutionary analysis of *yicC* orthologs by Maximum Likelihood method. The 51 *yicC* ortholog sequences were obtained from Ensembl Bacteria and their taxonomy information was extracted from NCBI Taxonomy. The evolutionary history was inferred by using the Maximum Likelihood method and a JTT matrix-based model (6) where the tree with the highest log likelihood is shown. The analyses were conducted in MEGA X (7, 8). Proteins from the bacteria of the same class are in the same color. The *E. coli* MG1655 *yicC* gene is highlighted in red, and a schematic of its protein domains is shown at the bottom.

**B**. The *sodB* fusion strains (WT: JC1248; *rne131*: JC1262; *rnc14*: JC1378) transformed with the pQE-80L vector or pYicC, expressing His-YicC, were induced with IPTG (50 μM) for 6 h at 37°C before measuring mCherry fluorescence.
**C**. The *sodB* fusion strains (WT: JC1248; *rnc14*: JC1378) transformed with the pQE-80L vector or pYicC were grown in LB+Amp at 37°C to an $OD_{600}$ of ~0.2, and then induced with 100 μM IPTG for 1 h. Total RNAs were prepared and analyzed by Northern blotting for RyhB and ChiX. SsrA served as an RNA loading control.
**D**. Coomassie staining of total protein samples prepared from WT (XL111) or Δ*pnp* (XL97) transformed with empty vector (pBR*) or pYicC2 and induced, as in **Figs. 7B, 7C,** resolved in a NuPAGE 4-12% Bis-Tris protein gel. Induced YicC proteins are underlined with red lines.
**E**. Northern blot comparing effect of YicC in WT and Δ*pnp* strains, quantitated in **Figs**. **7B, 7C**, and **S12F**. Asterisks indicate RyhB degradation products seen in Δ*pnp* cells.
**F**. Relative expression of sRNA MicA, ArcZ, and GcvB from **Fig. S12E** Northern blot. Results were plotted using data from two independent experiments.
Unpaired two-tailed Student's t-test was used to calculate statistical significance. Not significant (ns), $P > 0.05$; (∗) $P < 0.05$; (∗∗) $P < 0.01$.

### Supplementary Materials and Methods

#### Plasmid construction

All plasmids and DNA oligonucleotides used in this study are listed in Table S2 and S3. Plasmids deleted of candidate genes derived from the original positive clones were constructed using the QuikChange method with the following primers: JC214/215 (pΔ*cyaR*_glib), JC216/217 (pΔ*chiX*_glib), JC218/219 (pΔ*mcaS*_glib), JC220/221 (pΔ*rnt*_glib), JC242/243 (pΔ*yicC*_glib), JC248/249 (pΔ*yhbTS*_glib), JC226/227 (pΔ*aspX*_glib), JC222/223 (pΔ*rph*_glib), JC244/245 (pΔ*dinD*_glib), JC224/225 (pΔ*yhbQ*_glib) and JC246/247 (pΔ*yhbP*_glib). Gene deletions were verified by PCR with primers specific for gene flanking sequences.

pRnt, pYhbS and pYicC were constructed by standard restriction cloning that amplified the *rnt, yhbS* and *yicC* genes from *E. coli* MG1655 genomic DNA (gDNA) with JC208/209, JC268/269 and JC264/265 respectively, digested with BamHI/HindIII and then cloned into the pQE-80L plasmid (AmpR, Qiagen). This plasmid provides an N-terminal His tag, and thus these plasmids express His-tagged proteins. RNase T variant plasmids were constructed by the QuikChange method using pRnt as template with the following primers: JC296/297 (pRnt_R13A), JC298/299 (pRnt_D23A). pYicC2, and pAspXL, pAspXS, pAspXP plasmids were constructed by amplifying the *yicC* (5'UTR [-28 from ATG] and ORF) and *aspXL, aspXS, aspXP* genes from *E. coli* MG1655 genomic DNA (gDNA) with JC304/306 and JC270/277, JC272/277, JC273/277 respectively, and cloned into the pBR* plasmid (KanR, (9)), using restriction cloning (AatII/EcoRI). pAspXL_$m_{GCGC}$ and pAspXL_$m_C$ were generated by the QuikChange method using pAspXL as template with primers JC290/291 and JC288/289 respectively. XL1 blue chemical competent cells (Agilent) were used for cloning.

The BACTH plasmids pKT25-*yicC*, pKT25-*pnp*, pUT18-*yicC* and pUT18-*pnp* were constructed by restriction site cloning. Oligos JCO98/99, JCO98/101 and JCO96/97, JCO96/100 were used to PCR amplify *yicC* and *pnp* coding sequences, which then were digested with EcoRI/BamHI and cloned into pKT25 and pUT18 respectively. All plasmid clones were verified by Sanger sequencing (GeneWiz).

#### Strain construction

Strains, DNA oligonucleotides and gene fragments used in this study are listed in Table S1 and S3. Strains of fluorescent reporter fusions were constructed by λRed homologous recombineering in JC1183 ($Kan^R$) or JC1283 ($Kan^S$) of PCR products. To construct JC1183, *Cp26-cat-sacB-mCherry-sB* and *sB-R148K-sfGFP-T1-`lacY* were PCR amplified using primers JC148/145 and JC146/147 from JC1164 gDNA and pXG-30sf plasmid (10) respectively; the resultant PCR fragments were joined together by overlapping PCR using primers JC148/147; the full-length product was then recombineered in JC1095 (pSIM6) cells and selected on LB Chloramphenicol (10 μg/ml) agar plates. Recombination was verified by PCR using primers JC146/115, and the resulting strains were further tested for sucrose sensitivity on agar. To generate JC1283, the pCP20 plasmid (11) encoding the FLP recombinase was introduced into JC1183 to flip out the *kan* marker linked to the *CP18-araE*; this constitutive expression of AraE allows more linear induction from an arabinose promoter (12). To generate strain JC1164, PM1805 was P1 transduced with the *kan-*$P_{CP18}$*-araE* from BW27750 (12), and selected on LB+Kan agar. Then a PCR product of *pBAD-RBS1-mCherry-2xdBroccoli-T7t-`lacY* generated from overlapping PCR, using oligos JC125/122 (pET28c-F30-2xdBroccoli [a gift from Samie Jaffrey (Addgene plasmid # 66843 ; http://n2t.net/addgene:66843 ; RRID: Addgene_ 66843)] as template) and JC123/124 (pRSETb-mCherry (13) as template) for 1st-round PCR and oligo JC126/122 for 2nd-round PCR, was recombineered in the resultant strain, and selected on minimal glycerol sucrose agar; lastly, the PCR product *pBAD'-cat-sacB-'mCherry* amplified using oligos JC127/128 from PM1805 was recombineered in the resultant strain above and selected on LB+Chl10 agar. All chromosomal constructs were verified by PCR and the parental reporter fusion was further validated via Sanger sequencing of PCR products. All strains containing the *cat-sacB* cassette (for instance, JC1183) are sensitive to growth on sucrose and thus were used as recipients for recombineering genes of interest into fusions, selecting on minimal sucrose plates. Loss of chloramphenicol resistance was then monitored to confirm replacement of the cassette with the gene of interest.

The fluorescent reporter strains of *rpoS* (JC1197), *ompF* (JC1198), *chiP* (JC1200), *ompX* (JC1285), *mutS* (JC1286) were constructed by recombineering in JC1183 of PCR products amplified from MG1655 gDNA with JC118/179, JC180/174, JC182/178, JC278/279 and JC274/275 respectively, and selected on M63 glycerol sucrose agar. The formula for selection agar: for every liter, mix 490 mL 3% agar, 500 mL 2xM63 medium, 10 mL 20% glycerol, 50g sucrose, 1 mL 1% vitamin B1 and 300 $\mu$L 0.3% biotin. Reporter strains *RBS1* (JC1245), *sodB* (JC1246), *sdhC* (JC1334), *fumA* (JC1335) and *cirA* (JC1336) were constructed by recombineering in JC1283 of PCR products amplified from MG1655 gDNA with JC171/172, JC181/176, JC349/350, JC351/352 and JC353/354 respectively, and selected on the M63 glycerol sucrose agar. All gene reporter fusions were verified by PCR and sequencing using primers lacI-f/JC129.

JCM1001 and JC1316 were constructed by recombineering in NM541 of PCR products amplified from gDNA (*ybeW::zeo*) with JC36/37 and JC335/336, and selected on LB+Zeo agar. JC1318, JC1319 and JC1358 were constructed by PCR amplification from gDNA (*ybeW::zeo*) with primers JC337/338, JC341/342 and JC345/346 respectively and recombineered in JC1249 (pSIM6) and JC1246 (pSIM6) respectively; JC1262 was constructed by PCR amplification from gDNA (*ybeW::zeo*) with primers JC260/261 and recombineered in JC1248 (pSIM6); recombinants were selected on LB+Zeo agar. JC1327 was constructed by recombineering the gblock JCg20 in NM580 and selected on LB+1% arabinose agar via induction of a pBAD-*ccdB* toxin expression. $\Delta$*fur::cat* was introduced into the resultant strain via P1 transduction, using the P1 lysate from JC1249; transductants were selected on LB+Chl25. JC1282 was constructed by recombineering into JC1248 a PCR product amplified using JC292/293 from PM1805 and selected on LB+Chl10; the resultant strain was tested for sucrose sensitivity and then transformed with the pSIM6 plasmid. All chromosomal constructs were verified by PCR and the gene reporter fusions were further validated via Sanger sequencing of PCR products.

The chromosomal *ryhB* variant strains JC1291, JC1292 and JC1293 were constructed by recombineering in JC1282 using gBlock JCg10, JCg11 and JCg12 respectively, and recombinants were selected on M63 glycerol sucrose agar. *ryhB* chromosomal mutants were verified by PCR and sequencing with primers JC294/295. The chromosomal *aspX* variants JC1374 and JC1376 were constructed by recombineering gBlocks JCg23 in JC1246 (pSIM6) and JC1249 (pSIM6) respectively. JC1375 and JC1377 were constructed by recombineering gBlocks JCg24 in JC1246 (pSIM6) and JC1249 (pSIM6) respectively. Recombinants were selected on LB+Zeo agar, and recombination was verified by PCR amplification using primers JC240/241 followed by sequencing using primer JC240. Chromosomal gene mutations linked to an antibiotic marker were moved by P1 *vir* transduction as needed.

**5' Rapid Amplification of cDNA Ends (5'RACE)**

5' ends of AspX were mapped by 5'RACE as previously described (14). Briefly, total RNA of MG1655 grown in MOPS glucose (Teknova, M2106) to an $OD_{600}$ of ~1.0 was extracted. RNA samples were treated with BaselineZERO DNase (Epicentre Technologies) to remove DNA contaminants, then treated with Tobacco Acid Pyrophosphatase (TAP, Epicentre Technologies) to generate RNAs with a 5' monophosphate. RNA not treated with TAP (TAP -) was also included. After depletion of rRNA with Ribo-Zero kit (Illumina), an RNA adapter (FDM30: rArCrUrCrUrCrUrArCrUrGrUrUrUrCrUrCrCrArU) was then ligated to the 5'end of RNA with T4 RNA ligase 1 (New England Biolabs), and the resultant RNAs were reverse transcribed using SuperScript IV Reverse Transcriptase (Invitrogen) following standard protocols. After that, template RNA was digested by RNase H treatment (New England Biolabs), and cDNAs were amplified by PCR using the adapter-specific primer FDM32 (ACTCTCTACTGTTTCTCCAT) and an *aspX*-specific primer FDM110 (TGTACTACCCTGTACGAttaCTG). The PCR product was then TOPO cloned into the pCR4-TOPO vector (Invitrogen) and plasmids from transformants were sequenced and analyzed as previously described (15).

**Table S1 Bacterial strains used in this study**

| Strains | Description | Reference / Source |
|---|---|---|
| MG1655 | Wild type *E. coli K-12* strain | Lab strain collection |
| AZ234 | MC4100 Δ*hfq::cat-sacB* | (16) |
| BTH101 | *F-, cya-99, araD139, galE15, galK16, rpsL1 (Str r), hsdR2, mcrA1, mcrB1* | Euromedex |
| EM1238 | EM1055 Δ*ryhB::cat* | (17) |
| EM1320 | *rnc-14*::ΔTn10 | (18) |
| GSO145 | MG1655 Δ*cyaR::kan* | (19) |
| MG1054 | DJ480 Δ*fur::kan* | (16) |
| MG1056 | DJ480 Δfur::cat | Maude Guillier lab strain |
| MPK0106 | MG1655 Δ*abgR-ydaL::kan* | (20) |
| NM525 | MG1655 *lacIQ* FLP-scar | Lab strain collection |
| NM541 | MG1655 *lacIQ* FLP-scar *miniλ-tetR* | Lab strain collection |
| NM580 | MG1655 *lacIQ* FLP-scar *miniλ-tetR*, *lacI-T1T2-invZeoR-Kan-pBAD-ccdB-lacZ* for making transcriptional fusion | Lab strain collection |
| PM1805 | MG1655 *mal::lacIQ ΔaraBAD araC+ lacI'::pBAD-cat-sacB-lacZ miniλ-tetR* | (21) |
| XL1-blue | *recA1 endA1 gyrA96 thi-1 hsdR17 supE44 relA1 lac [F´ proAB lacIQ ZΔM15 Tn10 (Tetr )]* | Agilent Technologies |
| XL80 | NM525 Δ*pnp* | NM525 + P1 (Keio:JW5851) + pCP20 |
| XL97 | NM525 Δ*fur::cat* Δ*pnp* | XL80 + P1 (MG1056) |
| XL111 | NM525 Δ*fur::cat* | NM525 + P1 (MG1056) |
| JC1095 | MG1655 *mal::lacIQ ΔaraBAD araC+ lacI'::PBAD-recA-lacZ CP18-araE::kan* | (22) |
| JC1164 | MG1655 *mal::lacIQ ΔaraBAD araC+ CP18-araE::kan lacI'::pBAD-cat-sacB-mCherry-F30-2xdBroccoli-T7t* | This study[1] |
| JC1183 | MG1655 *mal::lacIQ ΔaraBAD araC+ CP18-araE::kan lacI'::CP26-**cat-sacB**-mCherry-R148K-sfGFP-T1, pSIM6* | This study[1] |
| JC1197 | MG1655 *mal::lacIQ ΔaraBAD araC+ CP18-araE::kan lacI'::CP26-**rpoS**-mCherry-R148K-sfGFP-T1* | This study[1] |
| JC1198 | MG1655 *mal::lacIQ ΔaraBAD araC+ CP18-araE::kan lacI'::CP26-**ompF**-mCherry-R148K-sfGFP-T1* | This study[1] |
| JC1200 | MG1655 *mal::lacIQ ΔaraBAD araC+ CP18-araE::kan lacI'::CP26-**chiP**-mCherry-R148K-sfGFP-T1* | This study[1] |
| JC1244 | JC1200 Δ*chiX::zeo* | JC1200 + P1 (JCM1001) |
| JC1245 | MG1655 *mal::lacIQ ΔaraBAD araC+ CP18-araE lacI'::CP26-**RBS1**-mCherry-R148K-sfGFP-T1* | This study[1] |
| JC1246 | MG1655 *mal::lacIQ ΔaraBAD araC+ CP18-araE lacI'::CP26-**sodB**-mCherry-R148K-sfGFP-T1* | This study[1] |
| JC1247 | MG1655 *mal::lacIQ ΔaraBAD araC+ CP18-araE lacI'::CP26-**chiP**-mCherry-R148K-sfGFP-T1* | JC1200 + pCP20 |
| JC1248 | JC1246 Δ*fur::kan* | JC1246 + P1 (MG1054) |
| JC1249 | JC1246 Δ*fur::cat* | JC1246 + P1 (MG1056) |
| JC1252 | JC1246 Δ*fur::kan* Δ*ryhB::cat* | JC1248 + P1 (EM1238) |
| JC1262 | JC1248 Δ*rne131::zeo* | This study[1] |
| JC1263 | JC1245 Δ*hfq::cat-sacB* | JC1245 + P1 (AZ234) |
| JC1264 | JC1197 Δ*hfq::cat-sacB* | JC1197 + P1 (AZ234) |
| JC1265 | JC1198 Δ*hfq::cat-sacB* | JC1198 + P1 (AZ234) |
| JC1269 | JC1248 Δ*hfq::cat-sacB* | JC1248 + P1 (AZ234) |
| JC1270 | JC1246 Δ*ryhB::cat* | JC1246 + P1 (EM1238) |
| JC1282 | JC1248 Δ*ryhB::cat-sacB pSIM6* | This study[1] |
| JC1283 | MG1655 *mal::lacIQ ΔaraBAD araC+ CP18-araE lacI'::CP26-**cat-sacB**-mCherry-R148K-sfGFP-T1 pSIM6* | JC1183 + pCP20 |

| | | |
|---|---|---|
| JC1285 | MG1655 *mal::lacIQ ΔaraBAD araC+ CP18-araE::kan lacI'::CP26-**ompX**-mCherry-R148K-sfGFP-T1* | This study[1] |
| JC1286 | MG1655 *mal::lacIQ ΔaraBAD araC+ CP18-araE::kan lacI'::CP26-**mutS**-mCherry-R148K-sfGFP-T1* | This study[1] |
| JC1291 | JC1249 *ryhB*_WT | This study[1] |
| JC1292 | JC1249 *ryhB_gcgc* | This study[1] |
| JC1293 | JC1249 *ryhB_g* | This study[1] |
| JC1296 | JC1285 *Δhfq::cat-sacB* | JC1285 + P1 (AZ234) |
| JC1297 | JC1286 *Δhfq::cat-sacB* | JC1286 + P1 (AZ234) |
| JC1316 | NM525 *Δfur::zeo* | This study[1] |
| JC1318 | JC1249 *sodB-FLAG::zeo* | This study[1] |
| JC1319 | JC1249 *ΔaspX::zeo* | This study[1] |
| JC1322 | JC1246 *Δfur::zeo* | JC1246 + P1 (JC1316) |
| JC1323 | JC1246 *ΔryhB::cat Δfur::zeo* | JC1270 + P1 (JC1316) |
| JC1327 | NM580 *ryhBp-rbs-lacZ* Δ*fur::cat* | This study[1] |
| JC1329 | MG1655 *mal::lacIQ ΔaraBAD araC+ CP18-araE lacI'::CP26-**rpoS**-mCherry-R148K-sfGFP-T1* | JC1197 + pCP20 |
| JC1330 | MG1655 *mal::lacIQ ΔaraBAD araC+ CP18-araE lacI'::CP26-**ompF**-mCherry-R148K-sfGFP-T1* | JC1198 + pCP20 |
| JC1332 | MG1655 *mal::lacIQ ΔaraBAD araC+ CP18-araE lacI'::CP26-**mutS**-mCherry-R148K-sfGFP-T1* | JC1286 + pCP20 |
| JC1334 | MG1655 *mal::lacIQ ΔaraBAD araC+ CP18-araE lacI'::CP26-**sdhC**-mCherry-R148K-sfGFP-T1* | This study[1] |
| JC1335 | MG1655 *mal::lacIQ ΔaraBAD araC+ CP18-araE lacI'::CP26-**fumA**-mCherry-R148K-sfGFP-T1* | This study[1] |
| JC1336 | MG1655 *mal::lacIQ ΔaraBAD araC+ CP18-araE lacI'::CP26-**cirA**-mCherry-R148K-sfGFP-T1* | This study[1] |
| JC1339 | JC1334 *Δfur::zeo* | JC1334 + P1 (JC1316) |
| JC1340 | JC1335 *Δfur::zeo* | JC1335 + P1 (JC1316) |
| JC1341 | JC1336 *Δfur::zeo* | JC1336 + P1 (JC1316) |
| JC1352 | JC1341 *Δhfq::cat-sacB* | JC1341 + P1 (AZ234) |
| JC1355 | JC1246 *ΔchiX::zeo* | JC1246 + P1 (JCM1001) |
| JC1356 | JC1246 *ΔmcaS::kan* | JC1246 + P1 (MPK0106) |
| JC1357 | JC1246 *ΔcyaR::kan* | JC1246 + P1 (GSO145) |
| JC1358 | JC1246 *Δrnt::zeo* | This study[1] |
| JC1359 | JC1246 *ΔyhbS::kan* | JC1246 + P1 (Keio: JW3125) |
| JC1360 | JC1246 *ΔaspX::zeo* | JC1246 + P1 (JC1319) |
| JC1361 | JC1246 *ΔyicC::kan* | JC1246 + P1 (Keio: JW3619) |
| JC1367 | JC1355 *ΔryhB::cat* | JC1355 + P1 (EM1238) |
| JC1368 | JC1356 *ΔryhB::cat* | JC1356 + P1 (EM1238) |
| JC1369 | JC1357 *ΔryhB::cat* | JC1357 + P1 (EM1238) |
| JC1370 | JC1358 *ΔryhB::cat* | JC1358 + P1 (EM1238) |
| JC1371 | JC1359 *ΔryhB::cat* | JC1359 + P1 (EM1238) |
| JC1372 | JC1360 *ΔryhB::cat* | JC1360 + P1 (EM1238) |
| JC1373 | JC1361 *ΔryhB::cat* | JC1361 + P1 (EM1238) |
| JC1374 | JC1246 *aspX_WT::zeo* | This study[1] |
| JC1375 | JC1246 *aspX*_$m_{GCGC}$*::zeo* | This study[1] |
| JC1376 | JC1249 *aspX_WT::zeo* | This study[1] |
| JC1377 | JC1249 *aspX*_$m_{GCGC}$*::zeo* | This study[1] |

| | | |
|---|---|---|
| JC1378 | JC1248 *rnc-14::tet* | JC1248 + P1 (EM1320) |
| JCM1001 | NM525 ∆*chiX::zeo* | This study[1] |

[1]. Described in Supplementary Materials and Methods (Strain construction)

**Table S2 Plasmids used in this study**

| Plasmids | Description | Marker | Reference / Source |
|---|---|---|---|
| pHDB3 | pBR322 deleted for the HindIII-AvaI fragment; the EcoRI-HindIII fragment was replaced by the pUC18 polylinker | AmpR | (2) |
| genomic library | Sau3AI partially digested (1.5-5 kb) LE392 genome ligated into BamHI digested pHDB3 vector | AmpR | (2) |
| pCyaR_glib | genomic library plasmid #3 hosted *cyaR* gene | AmpR | Library clone |
| pChiX_glib | genomic library plasmid #1 hosted *chiX* gene | AmpR | Library clone |
| pMcaS_glib | genomic library plasmid #20 hosted *mcaS* gene | AmpR | Library clone |
| pRnt_glib | genomic library plasmid #27 hosted *rnt* gene | AmpR | Library clone |
| pYicC_glib | genomic library plasmid #18 hosted *yicC* gene | AmpR | Library clone |
| pYhbS_glib | genomic library plasmid #4 hosted *yhbS* gene | AmpR | Library clone |
| pAspX_glib | genomic library plasmid #28 hosted *aspX* gene | AmpR | Library clone |
| pΔ*cyaR*_glib | genomic library plasmid #3 deleted *cyaR* gene | AmpR | This study[2] |
| pΔ*chiX*_glib | genomic library plasmid #1 deleted *chiX* gene | AmpR | This study[2] |
| pΔ*mcaS*_glib | genomic library plasmid #20 deleted *mcaS* gene | AmpR | This study[2] |
| pΔ*rnt*_glib | genomic library plasmid #27 deleted *rnt* gene | AmpR | This study[2] |
| pΔ*yicC*_glib | genomic library plasmid #18 deleted *yicC* gene | AmpR | This study[2] |
| pΔ*yhbTS*_glib | genomic library plasmid #4 deleted *yhbTS* genes | AmpR | This study[2] |
| pΔ*aspX*_glib | genomic library plasmid #28 deleted *aspX* gene | AmpR | This study[2] |
| pΔ*rph*_glib | genomic library plasmid #18 deleted *rph* gene | AmpR | This study[2] |
| pΔ*dinD*_glib | genomic library plasmid #18 deleted *dinD* gene | AmpR | This study[2] |
| pΔ*yhbQ*_glib | genomic library plasmid #4 deleted *yhbQ* gene | AmpR | This study[2] |
| pΔ*yhbP*_glib | genomic library plasmid #4 deleted *yhbP* gene | AmpR | This study[2] |
| pSIM6 | miniλ recombineering plasmid, pSC101 origin, *repAts* | AmpR | (23) |
| pCP20 | temperature-sensitive origin of replication, encodes the FLP recombinase | AmpR | (11) |
| pBAD33 | *araBAD* promoter-based expression vector , pACYC184 origin | CmR | (24) |
| pHfq | Hfq coding region cloned in pBAD33 | CmR | (25) |
| pBR*plac* | Expression vector, *Plac* promoter, pBR322 origin | AmpR | (26) |
| pQE-80L | Qiagen *lacIQ* vector for expressing N-terminally 6xHis-tagged proteins, ColE1 origin | AmpR | Qiagen |
| pBR* | A derivative of the pBR322-derived pBR*plac* vector, replacing the ampicillin cassette with kanamycin resistance cassette | KanR | (9) |
| pCyaR | *cyaR* sRNA gene cloned into pBR*plac* | AmpR | (27) |
| pChiX | *chiX* sRNA gene cloned into pBR*plac* | AmpR | (27) |
| pMcaS | *mcaS* sRNA gene cloned into pBR*plac* | AmpR | (27) |
| pArcZ | *arcZ* sRNA gene cloned into pBR*plac* | AmpR | (27) |
| pMgrR | *mgrR* sRNA gene cloned into pBR*plac* | AmpR | (27) |
| pDsrA | *dsrA* sRNA gene cloned into pBR*plac* | AmpR | (27) |
| pRprA | *rprA* sRNA gene cloned into pBR*plac* | AmpR | (27) |
| pMicF | *micF* sRNA gene cloned into pBR*plac* | AmpR | (27) |
| pRyhB | *ryhB* sRNA gene cloned into pBR*plac* | AmpR | (27) |
| pRnt | *His-rnt* ORF in pQE-80L (BamHI/HindIII) | AmpR | Vector provides $His_6$ tag; this study[2] |
| pRnt_R13A | *His-rnt* mutant R13 (CGT) -> A (GCT) on pQE-80L | AmpR | This study[2] |
| pRnt_D23A | *His-rnt* mutant D23 (GAT) -> A (GCT) on pQE-80L | AmpR | This study[2] |

| | | | |
|---|---|---|---|
| pYhbS | *His-yhbS* in pQE-80L (BamHI/HindIII) | AmpR | Vector provides $His_6$ tag; this study[2] |
| pYicC | *His-yicC* in pQE-80L (BamHI/HindIII) | AmpR | Vector provides $His_6$ tag; this study[2] |
| pYicC2 | *yicC* 5'UTR and ORF cloned into pBR* (AatII/EcoRI) | KanR | This study[2] |
| pAspXL | Putative long primary *aspX* (278 nts) cloned into pBR* (AatII/EcoRI) | KanR | This study[2] |
| pAspXS | Short primary *aspX* (118 nts) cloned into pBR* (AatII/EcoRI) | KanR | This study[2] |
| pAspXP | Processed *aspX* (98 nts) cloned into pBR* (AatII/EcoRI) | KanR | This study[2] |
| pAspXL_mGCGC | *aspXL* mutant G173C, C174G, G180C, C181G on pBR* | KanR | This study[2] |
| pAspXL_mC | *aspXL* mutant C177G on pBR* | KanR | This study[2] |
| pKT25-*yicC* | *yicC* ORF cloned into pKT25 (BamHI/EcoRI) | KanR | This study[2] |
| pKT25-*pnp* | *pnp* ORF cloned into pKT25 (BamHI/EcoRI) | KanR | This study[2] |
| pUT18-*yicC* | *yicC* ORF cloned into pUT18 (BamHI/EcoRI) | AmpR | This study[2] |
| pUT18-*pnp* | *pnp* ORF cloned into pUT18 (BamHI/EcoRI) | AmpR | This study[2] |

[2]. Described in Supplementary Materials and Methods (Plasmid construction)

**Table S3 Oligo primers, probes and gBlocks used in this study**

| Name | Sequence (5' to 3') | Description |
|---|---|---|
| JC125 | GTACAAGTAGctctacgacaacctcttcacagccaatctcTTGCCATGTGTATGTGGGAG | JC1164 construction |
| JC126 | acctgacgctttttatcgcaactctctactgtttctccatggagggtggaagtagagaag | JC1164 construction |
| JC127 | acctgacgctttttatcgcaactctctactgtttctccatAATGAGACGTTGATCGGCAC | JC1164 construction |
| JC128 | CCTTGATGATGGCCATGTTATCCTCCTCGCCCTTGCTCACACTGTCCATATGCACAGATG | JC1164 construction |
| JC145 | TATTACCCCCTTAATGTTTTTGCTTTCTGTGTgagattggctgtgaagaggttgtcgtagagttaCTACTTGTACAGCTCGTCC | JC1183 (*cat-sacB*) construction |
| JC146 | ctctacgacaacctcttcacagccaatctcACACAGAAAGCAAAAACATTAAGGGGGTAATAATGAGCAAAGGAGAAGAACT | JC1183 (*cat-sacB*) construction |
| JC147 | TTACGCGAAATACGGGCAGACATGGCCTGCCCGGTTATTACGGCGGATTTGTCCTACTC | JC1183 (*cat-sacB*) construction |
| JC148 | AACCCCGCTTATTAAAAGCATTCTGTAACAAAGCGGGACCCATTCTACAGTTTATTCTTGACATTGCACTGTCCCCCTGGTATAATAACTATAATGAGACGTTGATCGGCAC | JC1183 (*cat-sacB*) construction |
| JC118 | CCTTGATGATGGCCATGTTATCCTCCTCGCCCTTGCTCACATCATGAACTTTCAGCGTAT | JC1197 (*rpoS* fusion) construction |
| JC179 | TATTCTTGACATTGCACTGTCCCCCTGGTATAATAACTATTTCTGAGTCTTCGGGTGAAC | JC1197 (*rpoS* fusion) construction |
| JC180 | TATTCTTGACATTGCACTGTCCCCCTGGTATAATAACTATAGACACATAAAGACACCAAA | JC1198 (*ompF* fusion) construction |
| JC174 | CCTTGATGATGGCCATGTTATCCTCCTCGCCCTTGCTCACGATCACTGCCAGAATATTGC | JC1198 (*ompF* fusion) construction |
| JC182 | TATTCTTGACATTGCACTGTCCCCCTGGTATAATAACTATGTAGTCAGCGAGACTTTTCT | JC1200 (*chiP* fusion) construction |
| JC178 | CCTTGATGATGGCCATGTTATCCTCCTCGCCCTTGCTCACAGCAGCATAACCGGACTGAA | JC1200 (*chiP* fusion) construction |
| JC171 | TATTCTTGACATTGCACTGTCC | JC1245 (RBS1 control) construction |
| JC172 | CCTTGATGATGGCCATGTTATCCTCCTCGCCCTTGCTCACCATAGTTACCTCCTTAAAAT | JC1245 (RBS1 control) construction |
| JC181 | TATTCTTGACATTGCACTGTCCCCCTGGTATAATAACTATATACGCACAATAAGGCTATT | JC1246 (*sodB* fusion) construction |
| JC176 | CCTTGATGATGGCCATGTTATCCTCCTCGCCCTTGCTCACATATGGTAGTGCAGGTAATT | JC1246 (*sodB* fusion) construction |
| JC278 | TATTCTTGACATTGCACTGTCCCCCTGGTATAATAACTATTTAGGACTTATTTGAATCAC | JC1285 (*ompX* fusion) construction |
| JC279 | CCTTGATGATGGCCATGTTATCCTCCTCGCCCTTGCTCACCAGTGCTGAAAGACATGCAA | JC1285 (*ompX* fusion) construction |
| JC274 | TATTCTTGACATTGCACTGTCCCCCTGGTATAATAACTATTGCGCCTTATGTGATTACAA | JC1286 (*mutS* fusion) construction |
| JC275 | CCTTGATGATGGCCATGTTATCCTCCTCGCCCTTGCTCACATGGGCGTCGAAATTTTCTA | JC1286 (*mutS* fusion) construction |
| JC349 | TATTCTTGACATTGCACTGTCCCCCTGGTATAATAACTATTCTCCGGAACACCCTGCAAT | JC1334 (*sdhC* fusion) construction |
| JC350 | CCTTGATGATGGCCATGTTATCCTCCTCGCCCTTGCTCACAGGTCTTTGTTTTTTCACATTTCTTATc | JC1334 (*sdhC* fusion) construction |
| JC351 | TATTCTTGACATTGCACTGTCCCCCTGGTATAATAACTATACTCGCTTTTAACAGGGCAA | JC1335 (*fumA* fusion) construction |
| JC352 | CCTTGATGATGGCCATGTTATCCTCCTCGCCCTTGCTCACAGCCTGATAATGAAAGGGTTTG | JC1335 (*fumA* fusion) construction |
| JC353 | TATTCTTGACATTGCACTGTCCCCCTGGTATAATAACTATGCAATCAAAAAAGGCTGACAAATC | JC1336 (*cirA* fusion) construction |
| JC354 | CCTTGATGATGGCCATGTTATCCTCCTCGCCCTTGCTCACGACCCGTACGAAAGGGTTC | JC1336 (*cirA* fusion) construction |
| Verification oligos | | |
| lacI-f | TACGTTGACACCATCGAATGG | fusion verification |

| | | |
|---|---|---|
| JC115 | GTACATAATGGATTTCCTTACGCG | fusion verification |
| JC129 | CTCGAACTCGTGGCCGTTC | fusion verification |
| JC240 | CACTGCTAACAAAGAAGTGTG | *aspX* mutant verification |
| JC241 | CTTCCTTGTTTTTAACAAGTTG | *aspX* mutant verification |
| JC294 | CTGCCTGATGGCCTCGTAATC | *ryhB* mutant verification |
| JC295 | GTGGTGCTTTTGCCGAAGC | *ryhB* mutant verification |
| Oligonucleotides for chromosomal gene deletion | | |
| JC36 | CTGATGGCGGTCCAAAAAAAAGAGTCATCTTGCCTAAGAGGTTGACAATTAATCATCGG | JCM1001 (*chiX::zeo*) construction |
| JC37 | GTTGCGCTaaaAAAATGGCCAATATCGCTATTGGCCCGTCTCAGTCCTGCTCCTCGGCC | JCM1001 (*chiX::zeo*) construction |
| JC260 | CCGCTTCTTCGGCGCACTGAAAGCGCTGTTCAGCGGTtaaGTTGACAATTAATCATCGGC | JC1262 (*rne131::zeo*) construction |
| JC261 | CCCTGGCAGTTACCAGGGCTTGATTACTTTGAGCTAATTATCAGTCCTGCTCCTCGGCC | JC1262 (*rne131::zeo*) construction |
| JC335 | CAGATTCCGCatgACTGATAACAATACCGCCCTAAAGAAAGTTGACAATTAATCATCGGC | JC1316 (Δ*fur::zeo*) construction |
| JC336 | GGCTTTTCTCGTTCAGGCTGGCttaTTTGCCTTCGTGCGCTCAGTCCTGCTCCTCGGCCA | JC1316 (Δ*fur::zeo*) construction |
| JC341 | TCTGTTGACTGAAGCGGAACTTGACGATATTTTCTCCGTAtaaGTTGACAATTAATCATCGGC | JC1319 (Δ*aspX::zeo*) construction |
| JC342 | TCATCTGACGTGCCTTTTTTATTTGTACTACCCTGTACGATCAGTCCTGCTCCTCGGCCA | JC1319 (Δ*aspX::zeo*) construction |
| JC345 | AATGCGCGCTGCAATTTATCCGTATTAAGAGAATCAGATGGTTGACAATTAATCATCGGC | JC1358 (Δ*rnt::zeo*) construction |
| JC346 | TGCCTGAATCATTGCATGTCATCAGGCATCGACTCGATTATCAGTCCTGCTCCTCGGCCA | JC1358 (Δ*rnt::zeo*) construction |
| Oligonucleotides for chromosomal knock-in | | |
| JC337 | GCTGGTGAACTGGGAATTCGTAGCGAAAAATCTCGCTGCAgactacaaagacgatgacgacaagTAAGTTGACAATTAATCATCGGC | JC1318 (*sodB-FLAG::zeo*) construction |
| JC338 | AGCGTAGCGCTTCAGGCAATGCTGCATTTGCCATCAGTTATCAGTCCTGCTCCTCGGCCA | JC1318 *(sodB-FLAG::zeo*) construction |
| Oligonucleotides for on-plasmid gene deletion | | |
| JC214 | AAGGTTATAGCCTGTGTAATCTCCCTTACACGGGCTTATT | pΔ*cyaR*_glib construction |
| JC215 | CACAGGCTATAACCTTAATAAATGCAGCTGTATGTGATCG | pΔ*cyaR*_glib construction |
| JC216 | GATTGAGCGAGGCCAATAGCGATATTGGCCATTTTTttaG | pΔ*chiX*_glib construction |
| JC217 | GCTATTGGCCTCGCTCAATCTCACCATCCTGCCAATACTC | pΔ*chiX*_glib construction |
| JC218 | AAAACCGTTAAGAGTCTGGCGGATGTCGACAGACTCTATT | pΔ*mcaS*_glib construction |
| JC219 | CCAGACTCTTAACGGTTTTAACCTTTAGTTGCCAATTTTC | pΔ*mcaS*_glib construction |
| JC220 | GAGAATCAGATGTAATCGAGTCGATGCCTGATGACATGC | pΔ*rnt*_glib construction |
| JC221 | CGACTCGATTACATCTGATTCTCTTAATACGGATAAATTG | pΔ*rnt*_glib construction |
| JC242 | GGCGTTACGAGttacatAGACGTTCCTGTTTATAAAAGG | pΔ*yicC*_glib construction |
| JC243 | CAGGAACGTCTatgtaaCTCGTAACGCCAATTCTTAC | pΔ*yicC*_glib construction |
| JC248 | CCCCGGAGATTAcacGAGTTTTACTCCCTGTTTCAAC | pΔ*yhbTS*_glib construction |
| JC249 | GTAAAACTCgtgTAATCTCCGGGGTTTGCAGACTG | pΔ*yhbTS*_glib construction |
| JC226 | GAAGGTTACGtaaGGCACGTCAGATGACGTGCCTTTTTTCTTGTG | pΔ*aspX*_glib construction |
| JC227 | TGACGTGCCttaCGTAACCTTCGCACACTTCTTTGTTAGCAG | pΔ*aspX*_glib construction |
| JC222 | GAGATTTCAATATGTAGCGACGCAGAAGGCGGCGCTGGC | pΔ*rph*_glib construction |
| JC223 | CTTCTGCGTCGCTACATATTGAAATCTCCGGCTTGAAAC | pΔ*rph*_glib construction |
| JC244 | CGTCAGCAAGCGAATTGGCGTTACGAGttaTTCGATGTTC | pΔ*dinD*_glib construction |
| JC245 | ACGCCAATTCGCTTGCTGACGATGAAGCATTTAAGCAAC | pΔ*dinD*_glib construction |

| | | |
|---|---|---|
| JC224 | TGCGACGCAGACGATGTTAAAAGCGATTGAAATGCTCGTG | pΔ*yhbQ*_glib construction |
| JC225 | ATCGCTTTTAACATCGTCTGCGTCGCATGTTAAGGTCAG | pΔ*yhbQ*_glib construction |
| JC246 | ACCGCGAGGCGTTACATTGCTGTTCTCTTTTATACTGTGG | pΔ*yhbP*_glib construction |
| JC247 | GTATAAAAGAGAACAGCAATGTAACGCCTCGCGGTTAAACG | pΔ*yhbP*_glib construction |
| oligonucleotides for cloning and mutation | | |
| JC208 | GCG ggatcc TCCGATAACGCTCAACTTACC | pRnt constrution |
| JC209 | GCG aagctt CACCTCTTCGGCGGCAGATAG | pRnt constrution |
| JC296 | GGTCTGTGCGACgcTTTTCGTGGTTTTTATCCTGTTGTG | pRnt_R13A construction |
| JC297 | GATAAAAACCACGAAAAgcGTCGCACAGACCGGTAAGTTG | pRnt_R13A construction |
| JC298 | GTTGTGATCGcTGTTGAAACAGCCGGATTTAACGC | pRnt_D23A construction |
| JC299 | GTTTCAACAgCGATCACAACAGGATAAAAACCAC | pRnt_D23A construction |
| JC268 | GCG ggatccCTAATTCGAGTAGAAATTCCCATTG | pYhbS construction |
| JC269 | GCG aagcttAAAGCGATTGAAATGCTCGT | pYhbS construction |
| JC264 | GCG ggatccATCCGCAGTATGACCGCC | pYicC construction |
| JC265 | GCGaagcttttaTTCGATGTTCTGAATCTGCTCG | pYicC construction |
| JC304 | GCG gacgtcACCTCTCCTTTTATAAACAGGAACG | pYicC2 construction |
| JC306 | GCG gaattcttaTTCGATGTTCTGAATCTGCTCG | pYicC2 construction |
| JC270 | GCG gacgtcACAACTCTATCGGTATCGTTAC | pAspXL construction |
| JC272 | GCG gacgtcACAGAATCTGATGCACCC | pAspXS construction |
| JC273 | GCG gacgtcCTTACAAAGCAAAACGCTATAC | pAspXP construction |
| JC277 | GCG gaattcCAAGTTGATATTAGATTGTTATTTTTAAGTTAC | pAspX construction |
| JC288 | CTGATGCACgCGGCTTACAAAGCAAAACGCTATAC | pAspXL_mC construction |
| JC289 | CTTTGTAAGCCGcGTGCATCAGATTCTGTACGGAGAAA | pAspXL_mC construction |
| JC290 | CTGATcgACCCGcgTTACAAAGCAAAACGCTATACTGATG | pAspXL_mGCGC construction |
| JC291 | CTTTGTAAcgCGGGTcgATCAGATTCTGTACGGAGAAAATATC | pAspXL_mGCGC construction |
| Oligonucleotides for BACTH cloning | | |
| JCO96 | gcgGGATCCCCTTAATCCGATCGTTCGTAAATTCC | pKT25-*pnp* construction |
| JCO97 | gcgGAATTCttaCTCGCCCTGTTCAGCAG | pKT25-*pnp* construction |
| JCO98 | gcgGGATCCCATCCGCAGTATGACCGCCTAC | pKT25-*yicC* construction |
| JCO99 | gcgGAATTCttaTTCGATGTTCTGAATCTGCTCGC | pKT25-*yicC* construction |
| JCO100 | gcgGAATTCTCGCCCTGTTCAGCAGC | pUT18-*pnp* construction |
| JCO101 | gcgGAATTCgaTTCGATGTTCTGAATCTGCTCG | pUT18-*yicC* construction |
| gene fragments | | |
| JCg10 | GTCCCTTTCAACATCATTGACTTTCAAATGCGAGTCAAATGCATTTTTTT GCAAAAAGTGTTGGACAAGTGCGAATGAGAATGATTATTATTGTCTCgcg ATCAGGAAGACCCTCGCGGAGAACCTGAAAGCACGACATTGCTCACAT TGCTTCCAGTATTACTTAGCCAGCCGGGTGCTGGCTtttTTTTTGATCTTT CGTTCTCAATTTATCCACGGGAGTGCTTGTGTTCGTTATGCGCACTCCA GTAGGAACCACGTCCGCTTTGCGCTAAGGTGTAAATAACCACC | JC1291 construction |
| JCg11 | GTCCCTTTCAACATCATTGACTTTCAAATGCGAGTCAAATGCATTTTTTT GCAAAAAGTGTTGGACAAGTGCGAATGAGAATGATTATTATTGTCTCgcg ATCAGGAAGACCCTCGCGGAGAACCTGAAAGCACGACATTGCTCACAT TGCTTCCAGTATTACTTAGCCAcgCGGGTcgTGGCTtttTTTTTGATCTTTC GTTCTCAATTTATCCACGGGAGTGCTTGTGTTCGTTATGCGCACTCCAG TAGGAACCACGTCCGCTTTGCGCTAAGGTGTAAATAACCACC | JC1292 construction |

| | | |
|---|---|---|
| JCg12 | GTCCCTTTCAACATCATTGACTTTCAAATGCGAGTCAAATGCATTTTTTT GCAAAAAGTGTTGGACAAGTGCGAATGAGAATGATTATTATTGTCTCgcg ATCAGGAAGACCCTCGCGGAGAACCTGAAAGCACGACATTGCTCACAT TGCTTCCAGTATTACTTAGCCAGCCGcGTGCTGGCTtttTTTTTGATCTTT CGTTCTCAATTTATCCACGGGAGTGCTTGTGTTCGTTATGCGCACTCCA GTAGGAACCACGTCCGCTTTGCGCTAAGGTGTAAATAACCACC | JC1293 construction |
| JCg20 | CGCGCACCCACACCCAGGCCAGGGTGTTGTCCGGCACCACCTGGTCC TGGACCGCGCTGATGAACAGGGTCACGTCGTCCGTCCCTTTCAACATC ATTGACTTTCAAATGCGAGTCAAATGCATTTTTTTGCAAAAAGTGTTGGA CAAGTGCGAATGAGAATGATTATTATTGTCTCacgGTTATAAAGGAGGTc agctatgACCATGATTACGGATTCACTGGCCGTCGTTTTACAACGTCGTGA CTGGGAAAACCCTGGCGTTACCCAACTTAATCG | JC1327 construction |
| JCg23 | CACTGCTAACAAAGAAGTGTGCGAAGGTTACGTTTACAACTCTATCGGT ATCGTTACTTACCTGAACCCGTTCATCGGTCACCACAACGGTGACATC GTGGGTAAAATCTGTGCCGAAACCGGTAAGAGTGTACGTGAAGTCGTT CTGGAACGCGGTCTGTTGACTGAAGCGGAACTTGACGATATTTTCTCC GTACAGAATCTGATGCACCCGGCTTACAAAGCAAAACGCTATACTGAT GAAAGCGAACAGtaaTCGTACAGGGTAGTACAAATAAAAAAGGCACGTC AGATGACGTGCCTTTTTTCTTGTGAGGTTGACAATTAATCATCGGCATA GTATATCGGCATAGTATAATACGACAAGGTGAGGAACTAAACCATGGC AAAACTGACTTCAGCAGTACCAGTCCTTACCGCCCGCGACGTGGCAGG AGCCGTGGAGTTTTGGACCGACCGCTTAGGATTCAGCCGTGATTTCGT TGAAGATGATTTCGCGGGCGTTGTTCGCGACGACGTTACGCTGTTCAT TTCTGCAGTACAGGATCAAGTTGTGCCGGATAATACATTAGCATGGGTC TGGGTACGTGGTTTGGACGAGTTATACGCCGAGTGGTCAGAGGTGGTT TCCACGAACTTCCGTGACGCCTCTGGCCCTGCTATGACCGAGATTGGG GAACAGCCCTGGGGCCGCGAGTTTGCGTTACGTGACCCGGCTGGAAA TTGCGTCCACTTCGTTGCCGAAGAGCAAGATTGACAGTAACTTAAAAAT AACAATCTAATATCAACTTGTTAAAAAACAAGGAAGGCTAATATGCTAGT TGTAGAACTCATCATAGTTTTGCTGGCGATCTTCTTGGGCGCCAGATTG GGG | JC1374/JC1376 construction |
| JCg24 | GTCTGTTGACTGAAGCGGAACTTGACGATATTTTCTCCGTACAGAATCT GATcgACCCGcgTTACAAAGCAAAACGCTATACTGATGAAAGCGAACAGt aaTCGTACAGGGTAGTACAAATAAAAAAGGCACGTCAGATGACGTGCCT TTTTTCTTGTGAGGTTGACAATTAATCATCGGCATAGTATATCGGCATA GTATAATACGACAAGGTGAGGAACTAAACCATGGCAAAACTGACTTCA GCAGTACCAGTCCTTACCGCCCGCGACGTGGCAGGAGCCGTGGAGTT TTGGACCGACCGCTTAGGATTCAGCCGTGATTTCGTTGAAGATGATTTC GCGGGCGTTGTTCGCGACGACGTTACGCTGTTCATTTCTGCAGTACAG GATCAAGTTGTGCCGGATAATACATTAGCATGGGTCTGGGTACGTGGT TTGGACGAGTTATACGCCGAGTGGTCAGAGGTGGTTTCCACGAACTTC CGTGACGCCTCTGGCCCTGCTATGACCGAGATTGGGGAACAGCCCTG GGGCCGCGAGTTTGCGTTACGTGACCCGGCTGGAAATTGCGTCCACTT CGTTGCCGAAGAGCAAGATTGACAGTAACTTAAAAATAACAATCTAATA TCAACTTGTTAAAAAACAAGGAAGGCTAATATGCTAGTTGTAGAACTCA TCATAGTTTTGCTGGCGATCTTCTTGGGCGCCAGATTGGGG | JC1375/JC1377 construction |
| Northern DNA probes (5'biotinylated) | | |
| *ryhB* | AAGTAATACTGGAAGCAATGTGAGCAATGTCGTGCTTTCAGGTTCTC | (27) |
| *cyaR* | TGGTTCCTGGTACAGCTAGCATTTTATGGGTTATG | (27) |
| *arcZ* | GGCTAGACCGGGGTGCGCGAATACTGCGCCAACACCAGGG | (27) |
| *chiX* | CATTTTTTTATTATTATGCCGTCACTTTAAGCGACGGTG | (27) |
| *mcaS* | CCAGACTCTACAGTACACACAGCAG | (28) |
| *micA* | CATCTCTGAATTCAGGGATGATGATAACAAATGCGCGTCTTt | (27) |
| *gcvB* | CCAGAACACGCATTCCGATAAAACTTTTCGTTCCGGCTCAGG | (27) |
| *ssrA* | CGCCACTAACAAACTAGCCTGATTAAGTTTTAACGCTTCA | (27) |

| | | |
|---|---|---|
| *aspX* | GTACTACCCTGTACGAttaCTGTTCGCTTT | This study |
| *sodB* | TGCGATAGTCGATGTAATAAGCGTGTTCCCAGACATCAAC | (17) |
| *sdhC* | GGGGAACCGGATGGTCTGTAGGTCCAGATTAACAGGTC | (18) |
